# Reconfigurations within resonating communities of brain regions following TMS reveal different scales of processing

**DOI:** 10.1101/500967

**Authors:** Javier O. Garcia, Arian Ashourvan, Steven M. Thurman, Ramesh Srinivasan, Danielle S. Bassett, Jean M. Vettel

## Abstract

1

An overarching goal of neuroscience research is to understand how heterogeneous neuronal ensembles cohere into networks of coordinated activity to support cognition. To investigate how local activity harmonizes with global signals, we measured electroencephalography (EEG) while single pulses of transcranial magnetic stimulation (TMS) perturbed occipital and parietal cortices. We estimate the rapid network reconfigurations in dynamic network communities within specific frequency bands of the EEG, and characterize two distinct features of network reconfiguration, *flexibility* and *allegiance*, among spatially distributed neural sources following TMS. Using distance from the stimulation site to infer local and global effects, we find that alpha activity (8-12Hz) reflects concurrent local and global effects on network dynamics. Pair-wise allegiance of brain regions to communities on average increased near the stimulation site, whereas TMS-induced changes to flexibility were generally invariant to distance and stimulation site. In contrast, communities within the beta (13-20Hz) band demonstrated a high level of spatial specificity, particularly within a cluster comprising paracentral areas. Together, these results suggest that focal magnetic neurostimulation to distinct cortical sites can help identify both local and global effects on brain network dynamics, and highlight fundamental differences in the manifestation of network reconfigurations within alpha and beta frequency bands.

**Author Summary:** TMS may be used to probe the causal link between local regional activity and global brain dynamics. Using simultaneous TMS-EEG and dynamic community detection, we introduce what we call “resonating clusters”, or frequency band-specific communities in the brain, as a way to index local and global processing. These resonating clusters within the alpha and beta bands brain display both global (or integrating) behavior and local specificity, highlighting fundamental differences in the manifestation of network reconfigurations.

## 3 Introduction

The brain is an intricate collection of heterogeneous areas (Alivisatos et al., 2012), and a central goal of neuroscientific research is to understand how the coordination of these different regions support cognition (Azevedo et al., 2009; Bressler and Menon, 2010; Gollo et al., 2017). One theoretical approach encapsulates the coordinated activity into a framework of scales, and research has examined how local regional activity harmonizes with global signals (Bressler and Kelso, 2001). *Local activity* refers to cortical or thalamocortical interactions and reflects the transient coordination of inhibitory and excitatory neighboring neurons, constrained by basic neurophysiological factors such as refractory limitations and synaptic rising (Fries et al., 2007). However, research has shown that this local neural activity can be modulated by global activity in the brain (for review, see Buzsáki and Draguhn, 2004; Buzsáki and Wang, 2012). *Global activity* arises from propagation delays in cortico-cortical fibers and reflects the dynamic interactions and synchronization among distal networks. This conceptual framework of local and global networks interacting in cognitive processes is critical to the interpretation of neurophysiological signals. Yet, how this activity coheres to manifest cognition is still an active area of study (Bressler and Kelso, 2001; Cocchi et al., 2017).

EEG affords a natural way to study the scales of processing by examining oscillatory dynamics in different frequency bands (Buzsáki and Draguhn, 2004; Canolty and Knight, 2010). Changes in power in high frequencies, such as beta and gamma, have been used to infer local dynamics arising from the synchronization of populations of neurons (Brunel and Wang, 2003; Geisler et al., 2005). Similarly, the emergent activity in slower EEG frequencies, ranging across delta, theta, and alpha, has been interpreted as global activity arising from long-distance coordination of synchronized neural firing in disparate brain regions (Brunel and Wang, 2003; Geisler et al., 2005); however, there are known examples of cross-frequency interactions, suggesting an interaction between oscillatory activity that challenges a strict local/global interpretation on frequency dynamics (Canolty and Knight, 2010). In addition, results from EEG studies have indicated the importance of both local and global activity, indexed by high- and low-frequency oscillations, for understanding variability in human behavior (Nunez et al., 2001; Buzsaki, 2006; Nunez and Srinivasan, 2006; Volberg et al., 2009). However, EEG provides only an inferential framework to study interactions across scales of neural activity. Advancements in neurostimulation paradigms provide an avenue to directly study the causal role of local changes in oscillatory dynamics on global dynamics (Pascual-Leone et al., 2000; Bergmann et al., 2016), a long-known property of neurostimulation (Ilmoniemi et al., 1997).

Transcranial magnetic stimulation (TMS) has been proposed as a method to probe the dynamic interplay between local processing and the consequent global interactions with more distal regions of the brain (Massimini et al., 2009; Romei et al., 2012). Traditionally, single pulse TMS is a technique used to induce a short controlled burst of activity in a predetermined local brain region, directly causing a change in the local dynamics (Pascual-Leone et al., 2000). Research has identified behavioral outcomes from local stimulation for patients in clinical settings and healthy individuals in experimental tasks. Local stimulation in patients can successfully determine stroke recovery (for review, see Auriat et al., 2015), mitigate severe affective disorders (e.g., Berman et al., 2000), and preserve motor and language functions in presurgical mapping (Eldaief et al., 2013). TMS has also been successfully employed to confirm the role of an individual brain region on task performance, ranging from sensory attention (Taylor and Thut, 2012; Romei et al., 2013; Herring et al., 2015) to working memory performance (Brunoni and Vanderhasselt, 2014; Rose et al., 2016). Extant experimental research has suggested that the brain alterations caused by TMS is not limited to local perturbations (Ilmoniemi et al., 1997; Sale et al., 2015). By pairing TMS with other imaging modalities, it provides an innovative approach to study connectivity relationships among disparate brain regions (Siebner et al., 2009; Cocchi et al., 2015; Mancini et al., 2017).

Multimodal studies have paired TMS with functional neuroimaging, such as fMRI (e.g., Bestmann et al., 2008; Bohning et al., 1999), EEG (e.g., Bortoletto et al., 2015; Garcia et al., 2011), and PET (e.g., Paus, 1998), and measured stimulation-induced responses in brain areas that are distal to the stimulation site, indicating that stimulation can induce transient coordination between local and global activity (Paus, 1998; Bestmann et al., 2008; Driver et al., 2009). Complementing these findings, computational models of neurodynamics have demonstrated that regional differences in structural connectivity may account for how local network activity that is induced from a focal TMS pulse propagates along cortico-cortical fibers to influence global network synchronization (Muldoon et al., 2016; Gollo et al., 2017), and this research is corroborated with neurostimulation research that shows a structure-function constraint to the local stimulation and subsequent global (de)synchronization (Amico et al., 2017). While both experimental and modeling work has suggested the importance of interacting networks, few studies have employed the rich set of tools of network science to understand the propagation of stimulation (Bortoletto et al., 2015). Network science not only provides a mathematical language to describe complex connectivity patterns resulting from stimulation, but previous research has also proposed a variety of summary metrics in which to characterize local and global connectivity in the brain (for review, see Garcia et al., 2018). In this study, we address this existing gap in the literature and employ a method recently developed in network science to study the interactions of local connectivity and global network dynamics.

We investigated network reconfigurations from resting state EEG following single pulses of transcranial magnetic stimulation (TMS) using a method from network science that reveals modular architecture in the brain (Bassett and Bullmore, 2006; Bullmore and Sporns, 2012; Ercsey-Ravasz et al., 2013). Participants received single pulses of TMS to occipital or parietal cortex, and we computed functional connectivity using EEG data for a two second epoch surrounding stimulation (−1 to 1). Our theoretical question focused on the comparison between stimulation to spatially-disparate, large lobes of the brain, investigating how stimulation influenced network dynamics following stimulation as indexed by the modular architecture of the functional connectivity patterns. Each module is composed of regions with synchronized activity that are thought to be dynamically linked for the purpose of cohesive processing (Sporns et al., 2004; Achard et al., 2006; Bassett and Bullmore, 2006). To index local and global activity, we investigate two frequency bands that probe brain dynamics across these scales. We separately characterize the modular architecture of resting-state EEG within the alpha band and within the beta band, from which we define *resonating communities,* or communities of brain regions restricted to a specific frequency band. This delineation was inspired by the theoretical proposal by Rosanova and colleagues (2009) that posits brain regions have primary resonant frequencies: resting state activity is dominated by alpha in the occipital cortex, while parietal activity is dominated by beta activity. Consequently, our analysis investigates whether network changes in these frequency bands differentiate the location of the stimulation.

Changes in functional network organization before and after stimulation were characterized using two metrics from network science: *module allegiance* and *network flexibility.* Allegiance estimates how often regions are functionally connected with other regions, capturing stable subnetworks in the community structure across time points. Flexibility, in contrast, reveals the extent to which a region frequently (and flexibly) changes its assignment across communities between time points. Thus, allegiance increases the resolution of community assignments and captures coordinated activity of each node with every other node in the brain whereas flexibility examines whether a brain region changes affiliations overall. These two metrics are uniquely suited to investigate the scale of processing effects of stimulation since allegiance captures the unique shifts between each pair while flexibility identifies whether a node shifts its community affiliation across time. Together, the two metrics reveal how a stimulated region influences network dynamics. Our analyses extend previous research that has found network flexibility successfully characterizes large-scale functional differences (e.g., Telesford et al., 2016) in executive function (Braun et al., 2015) and mood (Betzel et al., 2017). Allegiance, on the other hand, has been used to describe observed network dynamics on a finer scale, estimating alignment with a pre-defined functional architecture (Bassett et al., 2015) as well as identifying transitions among certain network configurations (Ashourvan et al., 2017). Across the set of network science metrics adapted for neuroscience application (for review, see Garcia et al., 2018), allegiance and flexibility are the best suited to identify changes in scales of processing.

Using these measures, we report substantial differences between the alpha and beta band communities. While the alpha network reveals a dynamic interplay of local and global connectivity, communities within the beta band display a spatial specificity across both metrics, which suggest a more *local* connectivity impact of stimulation. Together, these results show that focal TMS to distinct cortical sites can help identify both local and global effects on dynamic network configurations, and demonstrate fundamental differences in the manifestation of network effects in alpha and beta frequency bands.

## 4 Results

Here we studied the brain dynamics following single pulses of TMS to occipital and parietal cortex using recently developed approaches from network science. First, we reconstructed estimated neural sources on a volumetric brain mesh and then extracted time series for 68 brain regions (Figure 1). Using a two second epoch surrounding stimulation (−1 to 1), we computed functional connectivity between all region pairs using the debiased weighted phase lag index (dwPLI) that has shown robustness to noise (Vinck et al., 2011; Vindiola et al., 2014). Our analysis focused on connectivity in the alpha and beta frequency bands since these bands have been suggested as resonant frequencies within the stimulated regions, alpha in occipital and beta in parietal regions (Laufs et al., 2003; Rosanova et al., 2009).

**Figure 1:**
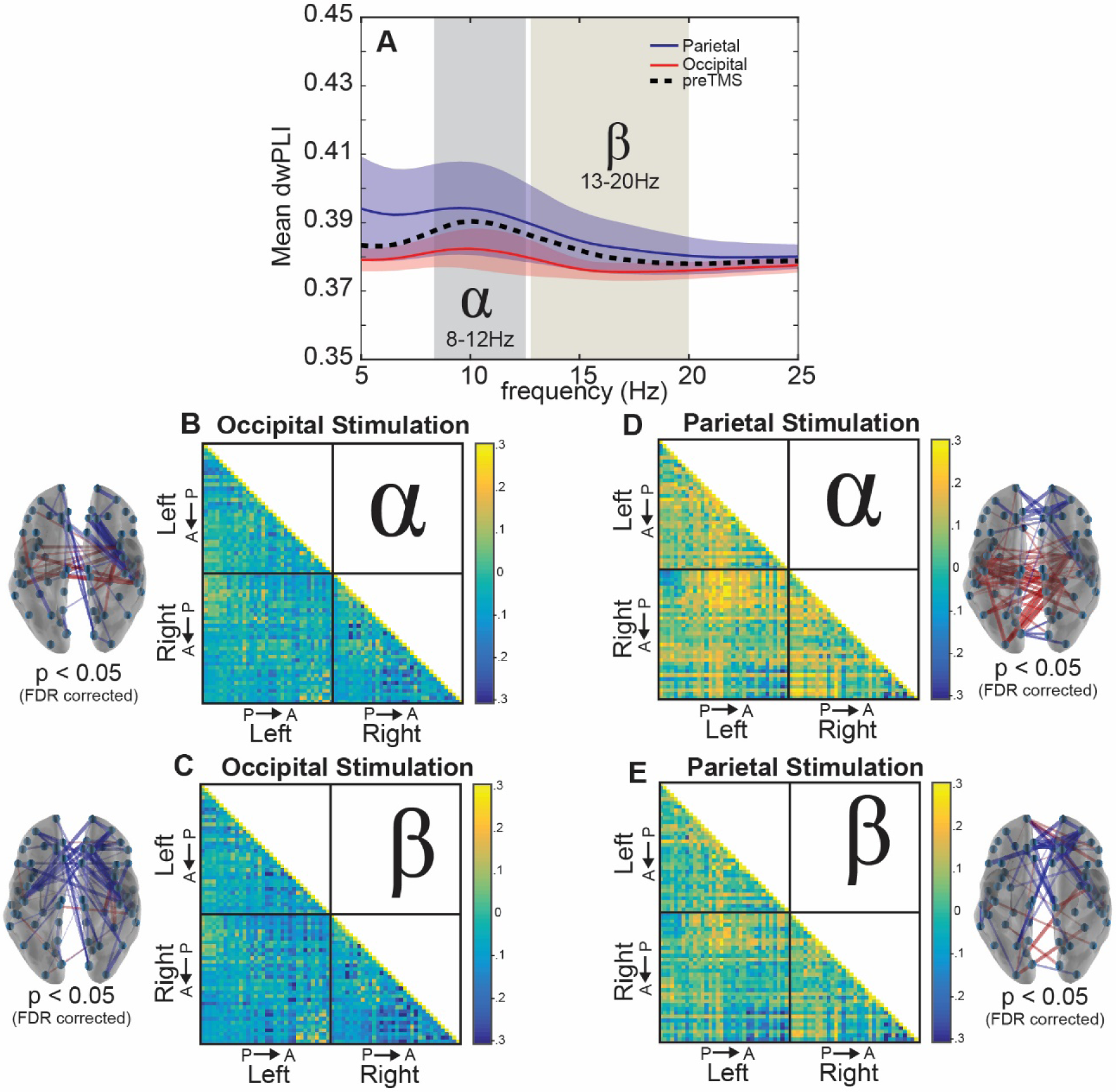
Whole-brain Connectivity Changes Following Stimulation. (A) Average dwPLI across the brain between 5Hz and 25Hz. (B, C) Debiased weighted phase lag index (dwPLI) differences between the second after TMS (postTMS) and the second before TMS (preTMS) intervals across trials averaged for occipital stimulation within the alpha band (B) and beta band (C). (D, E) Results similar to panels B-C but for parietal stimulation. Brain insets display the significant connections (p < .05, FDR adjusted) across the brain, providing a topographical illustration of the connectivity matrices where red lines indicate increased connectivity following stimulation and blue lines indicate decreased connectivity following stimulation.

### 4.1 Stimulation Effects on Whole-brain Connectivity

We began by examining patterns of functional connectivity in a whole-brain analysis (see Figure 1). We observed slightly higher connectivity across the brain within the alpha band (8-12Hz) both before (black dotted line in Figure 1A) and after stimulation to either site in bilateral occipital cortex (red line) or bilateral parietal cortex (blue line) compared to other frequency bands. This dominant response in whole-brain alpha synchrony likely reflects its role as a diffuse, communicative signal with multiple functions (Başar et al., 1999), serving as a global signal for sensory and information processing.

We next investigated changes in connectivity following stimulation by comparing changes between pre and post TMS intervals. As shown in Figure 1A, we observed that the average connectivity between all region pairs did not show much change within the alpha band after stimulation to either occipital or parietal sites (Occipital: t(9) = −0.95, p = 0.36; Parietal: t(9) = 1.05, p = 0.32, all uncorrected), and this was mirrored in the beta band with minimal connectivity differences for both stimulation locations (Occipital: t(9) = −1.39, p = 0.20; Parietal: t(9) = 0.41, p = 0.69, all uncorrected). However, there was a marked difference between occipital and parietal stimulation sites when examining the directionality and spatial specificity of the change following stimulation (Figure 1B-E). By subtracting the postTMS dwPLI estimate from the preTMS baseline, we observed a dispersed global decrease in connectivity for occipital stimulation (Figure 1B-C) for the regional pairs with the largest differences within the alpha and beta bands. Significant connections show some regional specificity, where the beta band shows decreases in connectivity between lateral central locations and medial frontal sites. The alpha band shows a similar connectivity pattern with an additional increase in connectivity between lateral regions toward the center of the brain. In contrast, we observed a marked increase within central and parietal sites as well as a frontal decrease in connectivity for parietal stimulation (Figure 1D-E). The alpha band shows a significant pattern of connectivity increases along in parietal regions, but this pattern is less robust within the beta band. Collectively, these whole-brain connectivity results show some frequency specificity for the stimulation sites, as might be predicted based on theories that suggest that stimulation could be facilitated or decremented by the inherent resonant frequency of the tissue (Rosanova et al., 2009). However, this observation could reflect the global influence of these regions on whole-brain connectivity rather than their targeted effects on subnetworks. Consequently, we next employed recent methods from network science to examine the effect of stimulation at a finer scale than average connectivity across nodes.

### 4.2 Community Organization in Resting Networks

To examine stimulation effects in brain communities, we capitalized on a network science approach that has been used previously to study modularity in brain networks. To estimate dynamic community structure, we optimize a multilayer modularity quality function, Q, using a Louvain-like greedy algorithm (Blondel et al., 2008; Mucha et al., 2010) to assign brain regions to communities, where each layer is a separate time slice. With this optimization, we extract our experimental *communities* by finding an optimal parameter scheme, which is composed of two parameters: (1) a structural resolution γ parameter and (2) a temporal resolution ω parameter. These two parameters determine the scale of the resulting graph, both structurally and temporally. As described in Garcia et al. (2018), there are several heuristics we may use to determine the optimal parameter for our dataset. We chose an unbiased “difference” heuristic because of the unique properties of this stimulation dataset. With this method, we compare the estimated Q from the preTMS interval to a Qnull derived from a shuffled null connectivity matrix where we shuffle the pair-wise dwPLI values, destroying the correlational structure observed in EEG data for each subject and parameter pairing. Each Q was then subtracted for each parameter pairing, comparing the observed model’s Q (from the unperturbed EEG connectivity patterns) and the null model’s Qnull (shuffled connectivity patterns) for each subject, and our analysis found a clear peak in the resulting Q matrix, suggesting that the range used was appropriate for this dataset.

This data-driven approach showed more local granularity in the network landscape following stimulation. Importantly, we defined network communities without stimulation during a period of rest. This allowed us to interrogate the dynamics of community reconfigurations following TMS, given a natural baseline, unbiased by the stimulation itself. Importantly, however, we interpret our results both within the confines of this community organization (Figures 3, 4) and outside of these confines (Figure 5). We defined network communities separately for both the alpha and beta bands, and used the most robust arrangement across the 100 iterations of modularity maximization as the final community structure. The 100 iterations of the preTMS interval were remarkably robust and consistent, showing 100% agreement across iterations for the alpha band and 98% agreement across the iterations within the beta band. We also observed noteworthy similarity (approximately 97% spatial similarity) between them except for a small cluster of motor-related brain regions (Figure 2).

**Figure 2:**
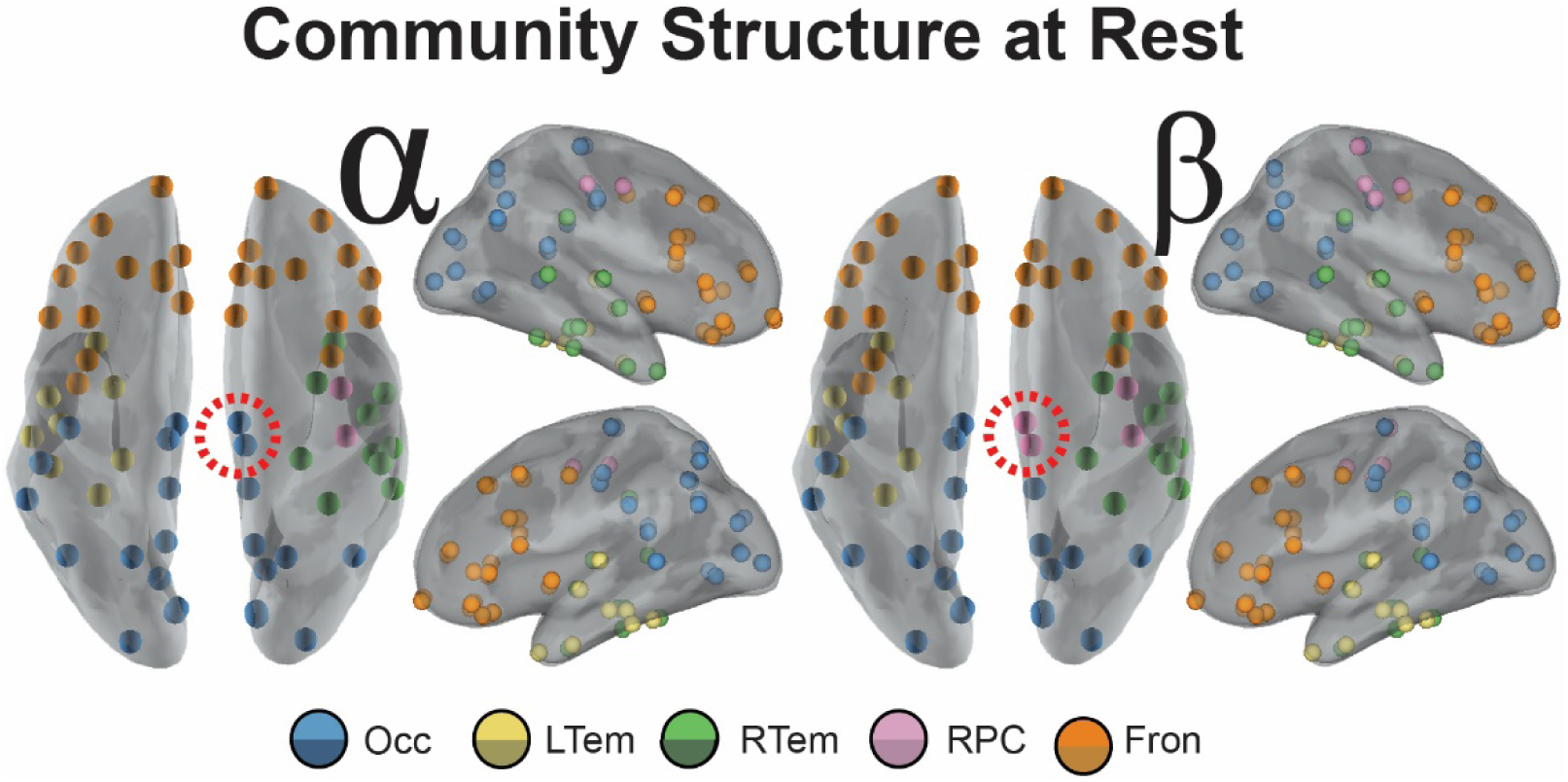
Communities derived from the interregional allegiance matrix in the preTMS interval for the alpha and beta bands. Inflated mesh visualizations of brain regions colored by community organization. Orbs are plotted at the centroid of the regions of interest. Community organization was found independently for the alpha band (Left) and beta band (Right) before stimulation with TMS. Dotted lines surrounding nodes near medial portion of the brains indicate the only two nodes unique to the different frequency bands.

**Figure 3:**
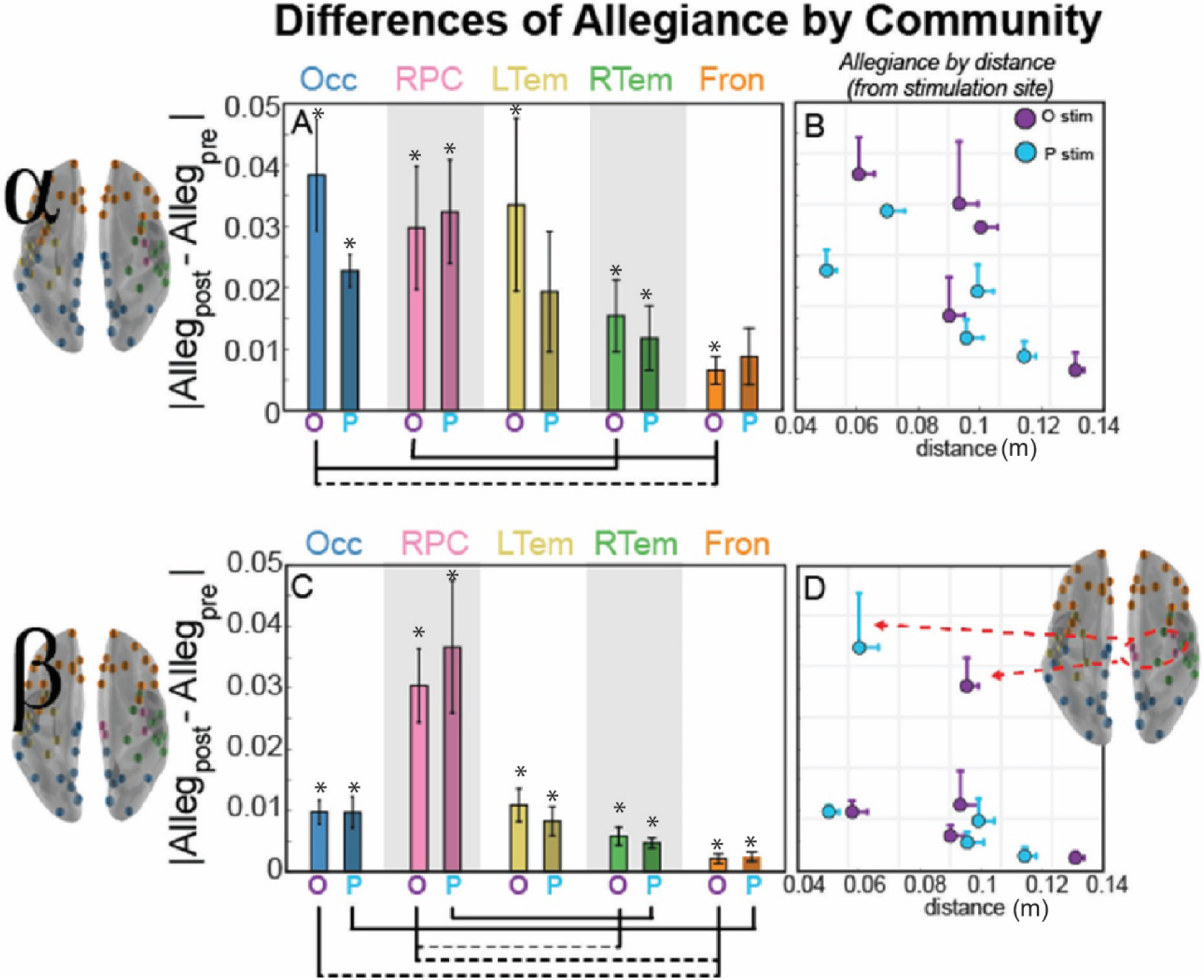
Community allegiance changes within alpha (top) and beta (bottom) band network as a function of distance from the stimulation site. (A, C) Bar plot of the mean magnitude change (SEM across subjects) from the preTMS interval in pair-wise allegiance from the stimulation site with the bar labeled O for occipital stimulation and P for parietal stimulation. For paired t-test between communities, dotted lines connecting communities indicate uncorrected significance (p < 0.05), while solid lines indicate significance corrected for multiple comparisons (Bonferroni, p < 0.05). (B,D) Scatter visualization of the mean magnitude allegiance change from the pre-TMS interval shown in (A,C), but now plotted as a function of distance from the stimulation site. Error bars indicate SEM across subjects (allegiance) or nodes within the community (distance) and the color of the marker indicates stimulation site (Occipital in purple and Parietal in blue). (*) indicate a significant difference from 0, indicating a change from the preTMS interval (p < 0.05, uncorrected). Brain inset for the beta band shows the nodes of the RPC community that are most affected by TMS regardless of stimulation site.

**Figure 4:**
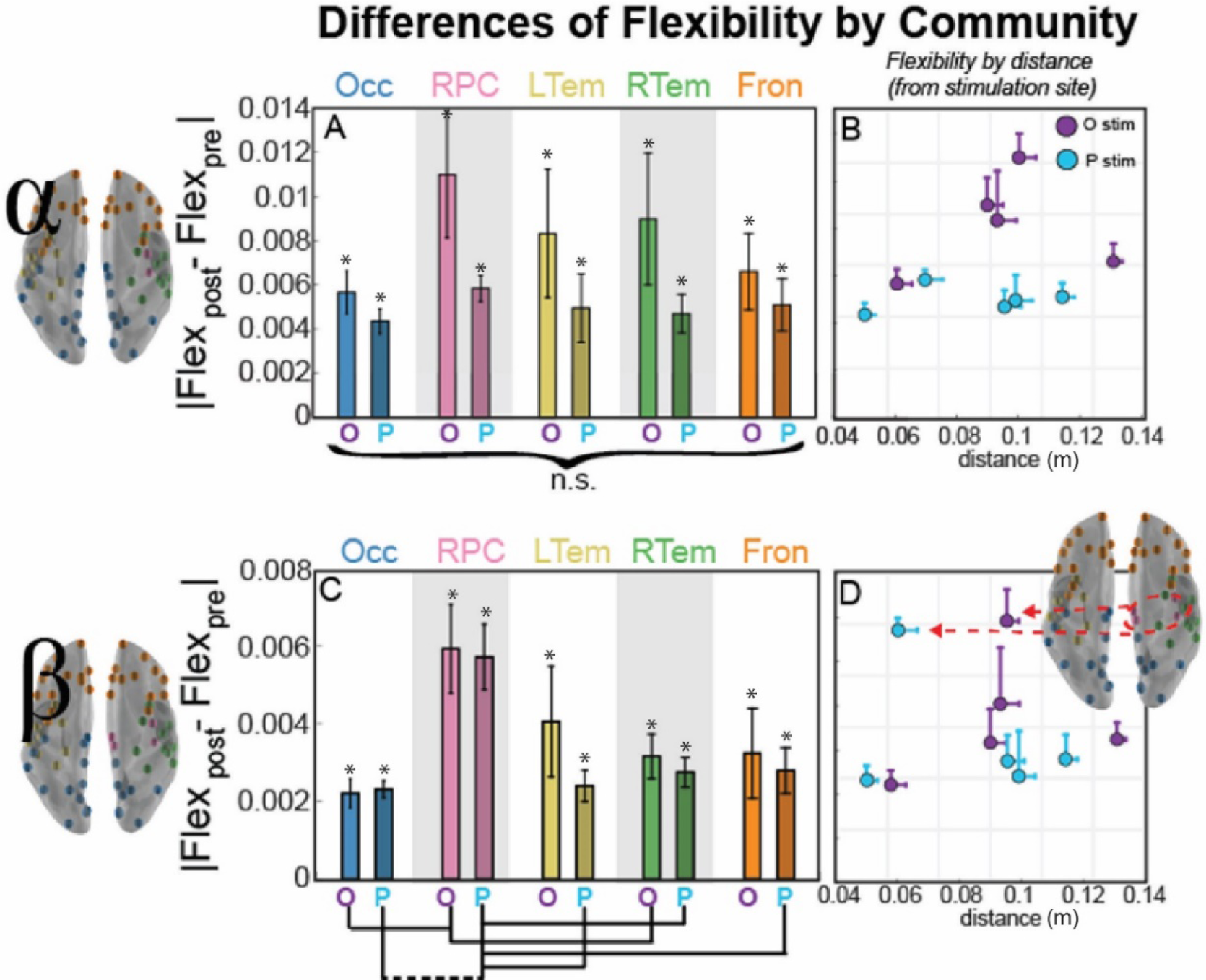
Community flexibility changes within alpha (top) and beta (bottom) band network as a function of distance from the stimulation site. (A, C) Bar plot of the mean magnitude change (SEM across subjects) from the preTMS interval in flexibility as a function of the distance from the stimulation site with the bar labeled O for occipital stimulation and P for parietal stimulation. For paired t-test between communities, dotted lines connecting communities indicate uncorrected significance (p < 0.05), while solid lines indicate significance corrected for multiple comparisons (Bonferroni, p < 0.05). (B,D) Scatterplot of the mean change in flexibility from the pre-TMS interval shown in (A,C), but now plotted as a function of distance from the stimulation site. Error bars indicate SEM across subjects (flexibility) or nodes within the community (distance) and the color of the marker indicates stimulation site (Occipital in purple and Parietal in blue). (*) indicate a significant difference from 0, indicating a change from the preTMS interval (p < 0.05, uncorrected). Brain inset shows the nodes of the RPC community that are most affected by TMS regardless of stimulation site.

**Figure 5:**
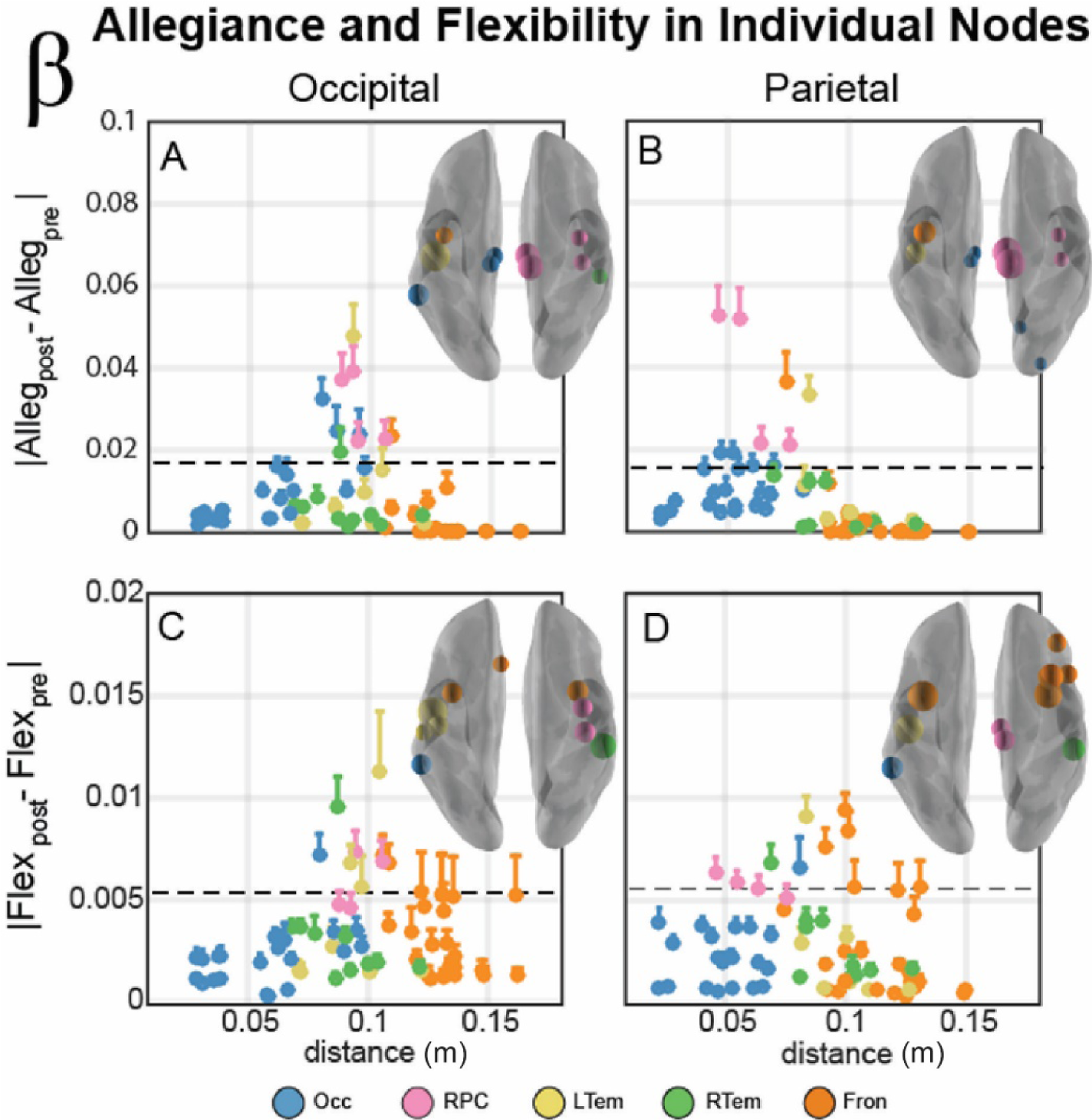
Individual node allegiance (A,B) and flexibility (C,D) changes within the beta band network. (A,B) Individual nodes module allegiance difference (Allegpost – Allegpre) plotted as a function of distance from the stimulation site for occipital (A) and parietal (B) stimulation sites. Color of node describes the community affiliation, and error bars indicate the standard error of the mean. Brain insets display the 85th percentile of module allegiance across all nodes, with nodes scaled by the relative magnitude of this allegiance change. The absolute magnitude of this percentile is also indicated by a horizontal dotted line in each plot. (C,D) Individual nodes flexibility difference (Flexpost – Flexpre) plotted as a function of distance from the stimulation site for occipital (A) and parietal (B) stimulation sites. Color of nodes, error bars, and brain inset display the same properties as above but in this panel with flexibility rather than allegiance.

In the alpha band (Figure 2, *left*), five communities are illustrated: a bilateral occipital-parietal network (Occ, blue), a right paracentral community (RPC, pink), a left temporal network (LTem, yellow), a right temporal network (RTem, green), and a bilateral frontal network (Fron, orange). This largest community (Occ) is a large cluster of regions in occipital and parietal cortex, an organization that is perhaps unsurprising, given the commonly observed peak of the alpha rhythm within occipital-parietal regions (Hari and Salmelin, 1997). Interestingly, five similar communities were also found within the beta band (Figure 2, *right*), and the only observed difference was in the right paracentral community (DKT atlas regions: R precentral, R postcentral). In addition to the two nodes of the pre- and post-central sulcus in the alpha RPC community, the beta band RPC community also contained regions of the medial paracentral lobule, putative sources of motor-related planning (DKT atlas regions: R paracentral, R posterior cingulate). This RPC community in beta nicely aligns with previous literature that implicates the beta band in motor-related activity (Pfurtscheller et al., 1996b), providing support that the detected network communities captured frequency-specific effects.

### 4.3 Community Allegiance Differentiates Beta Band Communities from the Alpha Band

We next sought to characterize how stimulation influenced dynamic network reconfigurations from the natural baseline resting state. We employed a metric of allegiance that captures how often two nodes are present within the same community before and after stimulation. Figure 3 (A, C) shows the average pair-wise difference in allegiance before and after stimulation within each of the five communities identified from the preTMS resting state connectivity (Alleg_post_-Alleg_pre_). Within the alpha band (top row), we observe some specificity to the stimulation site. In comparing allegiance for each pairing of the communities, allegiance change is highest for the occipital-parietal community (Occ) and lowest for the frontal community (Fron; paired t-test, t(9) = 3.8, p = 0.004, uncorrected), and this was true for each of the subjects within our sample (see *Supplemental Figure 6* to view more about robustness of effects across subjects). The beta band (bottom row), however, shows clear community specificity, where allegiance of the right paracentral (RPC) community is significantly higher than the right temporal (RTem) and frontal (Fron) communities following stimulation (paired t-tests; RTem, t(9) = −2.6, p = 0.028; Fron, t(9) = −2.8, p = 0.020, all uncorrected). To speak to robustness, the RPC showed the highest allegiance in 80% and 60% of subjects for occipital and parietal stimulation, respectively. Also, nearly each average change is significantly different from the preTMS resting state estimate (labeled with * in Figure 3B) with the exception of parietal stimulation effects in LTem and Fron.

We also examined whether the community allegiance depended on distance from the stimulation site, which we operationalized as the Euclidean distance from the centroid of the community to the node closest to the stimulation site, estimated in meters from a reconstructed 3D mesh. The effects of stimulation within the alpha band revealed that the nodes closest to the stimulation site are most susceptible to stimulation, and as Figure 3A shows, this effect is reduced for the communities further from the stimulation site (see *Supplemental Figure 7* for a non-parametric correlational analysis with distance and the graph metrics).

In contrast, the RPC community in the beta band was impacted more strongly by stimulation with high specificity (Figure 3D, pink RPC nodes) by comparison to the other communities. Thus, for the beta band, stimulation didn’t follow a simple rule based on distance from the stimulation site as observed in alpha; instead, the stimulation effect was strongest in the RPC community, suggestive of resonant frequencies within the regions targeted (Rosanova et al., 2009). This observation aligns with the *natural frequencies* account of stimulation based on the strong role that beta band serves in motor-related activity (Pfurtscheller et al., 1996b), and the prevalence of motor regions within the RPC community. These results indicate the importance of considering pairwise regional activity within a community, so we next examined a network measure of flexibility to investigate regional dynamics.

### 4.4 Flexibility Differences Indicate Whole-brain Effects within the Alpha Band

As a complement to allegiance, which measures the temporal consistency of community structure at the inter-regional level, we investigated flexibility, a metric that describes how often each node changes the community to which it is allied. This analysis captures whether stimulation drives certain brain regions to cohere with different communities in a manner that is different from their community participation prior to stimulation.

Figure 4 shows the differences in flexibility, averaged across nodes within a community, before and after stimulation (Flex_post_-Flex_pre_). First, we compared flexibility in these communities to 0, or no difference between Flex_post_ and Flex_pre_. We see that all communities have a significant change in flexibility in both the alpha and beta communities, suggesting a robust change in flexibility after stimulation reflecting the causal role of TMS pulses on dynamic network reconfigurations. Also, overall, we see a large difference in effect size for the different frequency bands, with alpha communities showing more flexibility overall.

We next compared each community pair to understand the specificity of these effects. Within the alpha band (top row), we observed minimal difference between communities or stimulation sites; rather, TMS is associated with a statistically equivalent change in flexibility across communities. For the beta band (bottom row), a single community stands out. The RPC community is again the most flexible following TMS. For occipital stimulation, flexibility of the right paracentral (RPC) community is significantly higher than the occipital-parietal (Occ) and trending for right temporal (RTem) communities (paired t-tests; Occ, t(9) = −3.0, p = 0.016, uncorrected; RTem, t(9) = −2.2, p = 0.057, uncorrected). This difference is even stronger for parietal stimulation, where flexibility for RPC is tends to be higher than that observed in the left temporal (LTem), right temporal (RTem), frontal (Fron), and significant (Bonferroni corrected) when compared to occipital-parietal (Occ) communities (paired t-tests; LTem, t(9) = −3.1, p = 0.013; RTem, t(9) = −2.6, p = 0.030; Fron, t(9) = −3.1, p = 0.014; Occ, t(9) = −4.0, p = 0.003, all uncorrected). These results are reminiscent of the pair-wise allegiance difference showing an increase within a single community (Figure 5C).

We next examined whether flexibility depended on distance from the stimulation site. Alpha communities showed minimal dependence between distance and flexibility (Figure 4B), but there was no significant difference across any of the communities (Figure 4A). In contrast, when we considered flexibility within the beta band, we observed that the RPC community displayed the strongest effect of stimulation (Figure 4D). Combined, these flexibility results demonstrate consistency with the allegiance results, suggesting an effect of resonant frequency in the RPC community for the beta band. However, the communities have variable numbers of nodes and a few nodes could substantially influence the means shown in Figures 4 and 5, so our final analysis examined individual node dynamics to determine whether the smaller size of the RPC community was the primary driver of beta band effects.

### 4.5 Individual Node Clusters Suggest a Reconfiguration of the Beta Band Network after TMS

Since all of the previous analyses examined only the overall community differences, our final analysis examined the individual node allegiance to the stimulation site and flexibility changes after stimulation. This analysis examines the spatial specificity of the TMS modification of the graph metrics that may be masked by averaging across many nodes within the affiliated community. Figure 5 displays individual node magnitude allegiance (top row) and flexibility (bottom row) differences in the 5 communities of the beta band network following stimulation to occipital (left) and parietal (right) cortex. Overall, there is a change in allegiance within a narrow range of distances from the stimulation site when considering both occipital (Figure 5A) and parietal stimulation (Figure 5B). Although regions in the right paracentral community (RPC, pink) show substantial change in allegiance, several nodes from other communities also have a similar response profile. To examine the spatial topology of these effects, the brain insets illustrate nodes corresponding to top 15% of allegiance changes (nodes on or above the threshold line in Figure 5A-B). The most influenced nodes surround the sensorimotor regions of the brain, including the RPC community and nearby regions around the central sulcus, the rolandic sites of the brain.

In contrast, the individual node flexibility changes are more diffuse (Figure 5, bottom row). The nodes corresponding to top 15% of flexibility changes are plotted above the threshold line, and these effects span a larger range of distances than the allegiance changes. However, the spatial topology is quite similar. Changes in flexibility after stimulation are strongest in a cluster of nodes surrounding the central sulcus. Collectively, these results reveal that the nodes in the RPC community were not uniquely influenced; instead, the network dynamics of the RPC as well as the surrounding bilateral sensorimotor regions showed the largest flexibility and allegiance changes within the beta band communities. Together these results suggest a rapid *reconfiguration* of the resting beta community organization following TMS stimulation, rather than enhancement of a single community.

## 5 Discussion

We investigated network reconfiguration in resting state EEG following single pulses of transcranial magnetic stimulation (TMS) using a method from network science that allows for a quantitative description of the brain’s modular architecture (Bassett and Bullmore, 2006; Bullmore and Sporns, 2012; Ercsey-Ravasz et al., 2013). Our analysis focused on connectivity differences between the one second before stimulation and the one second after stimulation. More specifically, we examined network differences in the alpha and beta frequency bands since these bands have been suggested as resonant frequencies within the stimulated regions: alpha in occipital and beta in parietal regions (Laufs et al., 2003; Rosanova et al., 2009).

Our results first examined whole-brain effects of stimulation, and we observed differential effects on connectivity between stimulation sites, although no stark differentiation was seen between the alpha and beta frequency bands. Next, we employed a network theoretical approach to identify communities from the resting state EEG data, and we observed several differences in the structure of functional connectivity in both frequency bands after TMS. Within the alpha band, stimulation produced local effects, but interestingly, stimulation also produced more global effects, altering network flexibility across *all occipital, parietal, paracentral, and frontal* communities when applied to either occipital or parietal cortex. In stark contrast, beta band activity showed high specificity of TMS-induced effects on allegiance and flexibility within a rather focal paracentral network near sensorimotor cortex. These novel results using network science approaches with TMS/EEG revealed an interesting interplay between local and global activity across frequency bands that might underlie how network reconfigurations give rise to coordinated brain activity.

### 5.1 Global effects within the alpha band manifest in similar flexibility across communities

We have shown that TMS to resonating communities constrained within the alpha band has equivalent impact on the overall flexibility of all communities. This finding implies a global impact of stimulation, regardless of stimulation site, to the alpha band networks. Since the first observations of the alpha band by Berger (1929), alpha band activity (8-12Hz), also known as the *Berger rhythm*, has been a brain rhythm of frequent study due to its dominance in resting EEG, and it is often the only visually observable pattern to the naked eye in the EEG trace. Since its first observation, several hypotheses have been proposed ascribing a functional role to its presence in EEG. The first theory was proposed by Adrian and Mathews (1934) who found that the power within the alpha band increases when subjects are awake with eyes closed. They interpreted this as alpha band activity reflecting a brain-state of inactivity, priming the brain for incoming information. This theory has been expanded and revised to more clearly represent ‘cortical idling’ (Pfurtscheller et al., 1996a). Recently, however, this theory has been somewhat abandoned because of the difficulty of reconciling it with behavioral experiments that indicate a functional role for power within this band. For example, alpha band activity is also associated with working memory load (Klimesch, 1996; Jensen et al., 2002; Tuladhar et al., 2007). Thus, beyond this ‘spontaneous’ alpha rhythm, researchers have discovered other forms of alpha, so-called ‘functional’ alpha (Başar, 2012), which is observed during many cognitive and motor processes. These theories have been further broadened, suggesting alpha activity may even be an *access controller* to a knowledge system (Klimesch, 2012). Collectively, however, alpha activity consistently represents a diffuse, communicative signal with multiple functions, an arguably global signal in terms of its impact on sensory information.

Research using TMS to investigate the functional connectivity within the alpha band, however, provides a more limited view of alpha activity. For example, Rosanova et al. (2009) showed that enhancement of the alpha band is primarily restricted to occipital cortex, regardless of stimulation site. These researchers noted that occipital cortex might *resonate* at the alpha frequency. Our results expand the theory proposed by Rosanova and colleagues by showing that the brain may be parsed into separate resonating communities within the alpha band and each cluster is overall equally modified by TMS, as indicated by the similar flexibility metrics across the resonating communities. Interestingly, however, nodal allegiance to the stimulation site reveals a rather direct and localized impact within the occipital and parietal cortex: specifically the cluster that is most associated with alpha band activity. Together, these results suggest a process whereby alpha connectivity can provide both the specific visual effects shown in early studies while also serving many functional roles across disparate brain regions.

### 5.2 Local effects of beta band manifest in the specificity of network changes within the paracentral community

Two pieces of evidence in the current study converge to show that the paracentral network plays a highly specific role in dynamic network reconfiguration within the beta band, and our results support and extend the natural frequency theory of the brain following single pulses of TMS (Rosanova et al., 2009). First, the network communities defined on resting state activity are identical between alpha and beta bands, except for the right paracentral network (RPC). The beta-band RPC consists of nodes within the paracentral lobule and two nodes within the pre and post central gyri that have been in sensorimotor processing, and previous research has consistently identified motor-related activity in the beta band (Pfurtscheller et al., 1996b). Second, following a single TMS pulse to either stimulation site, we observe that the paracentral region uniquely displays both the highest flexibility and allegiance changes from the baseline state in the beta band, but not the alpha band. Together, these findings suggest a unique specificity that would support the natural frequency theory of TMS and are aligned with other network metrics suggesting beta band influence after parietal stimulation (Amico et al., 2017). In fact, our results provide intuition at a level of granularity that has not been previously explored, by capitalizing on recently developed methods from network science (Bassett and Sporns, 2017) to investigate perturbations of brain networks following single pulses of TMS.

The granularity of this effect was further enhanced by inspection of the nodal allegiance and flexibility across the brain. The RPC community was clearly involved and appeared to be an isolated community with increased nodal allegiance (among its nodes) and global flexibility in the beta-band communities. However, when we inspected the single nodes contributing to this effect, we found that this effect was not constrained by the boundaries of the beta-band RPC network as we defined from baseline activity. Instead, the effects spread to nodes outside this RPC community and consisted of a cluster of nodes surrounding the central sulcus. This granularity of the beta band network effects after TMS aligns well with the well-known involvement of rolandic sites in sensorimotor processes and discharges of beta-band activity (Baker, 2007).

### 5.3 Clinical Implications

TMS has been used successfully in clinical settings for treating movement disorders (Pascual-Leone et al., 1994) and mitigating severe affective disorders (Berman et al., 2000); however, some studies that have investigated the efficacy and efficiency of TMS treatment for depression (for review, Loo & Mitchell, 2005) suggest that most treatment regimens are suboptimal, often stimulating for a duration of two weeks and only having a minor benefit. Here we have shown a complex interplay between local and global neural processing, but more generally, our results speak to the specificity of TMS, where particular resonating communities of brain regions (e.g., beta band activity emanating from sensorimotor regions) or diffuse sets of resonating communities (e.g., all regions in alpha band) may be modified by TMS regardless of stimulation site. In other words, we show that stimulating a focal region can have distal effects on many other brain regions. Future studies expanding on how individual variability in brain connectivity impacts how TMS propagates through cortex may eventually reveal the specific networks or brain regions that may predict successful treatment. We believe the methods and initial results within this work hold promise in future studies to help determine stimulation protocols for a variety of clinical settings and surrounding several cognitive domains.

Our results first examined whole-brain effects of stimulation, and we observed differential effects on connectivity in both alpha and beta activity, although no stark differentiation was seen between stimulation to occipital or parietal sites, globally. Next, we employed a network theoretical approach to identify communities from the resting state EEG data, and we observed several differences in the structure of functional connectivity in both frequency bands after TMS. Within the alpha band, stimulation produced local effects, but interestingly, stimulation also produced more global effects, altering network flexibility across *all* communities when applied to either occipital or parietal cortex.

Despite these general effects, our coarse-level results are merely the first step as much more must be completed to determine the robustness of much of these network dynamics that may include any gender differences (70% of this sample were male), individual susceptibility to stimulation, or state-based stimulation specificity (Thut et al., 2011), of which our current study does not tackle. We further expand on other methodological considerations that may guide future studies within this domain in the following.

### 5.4 Methodological considerations

Our use of community detection to understand functional connectivity in the brain and the effects of TMS on specific brain networks focuses on two stimulation sites and two common frequency bands. We use a phase-based undirected connectivity measurement and inspect graph metric changes at a single snapshot of the available parameters within the dynamic modularity framework we applied. This initial implementation of a graph theoretical approach on human neurostimulation effects may be expanded in the future to investigate the directed communication between these resonating communities (e.g., Reimann et al., 2017), cross-frequency communication (Canolty and Knight, 2010), and increased resolution across frequencies of the brain. The methodological choices within this work also focused on merely one spatiotemporal scale, which may not completely account for the spectral sensitivity across the regions of the brain, and the results target a wakeful resting state in indivduals, and it may not extend to the *active* or *sleeping* brain (Massimini et al., 2005; Hasson et al., 2009). Future research may extend this work to take into account the spectral macroarchitecture of evoked and induced oscillations in the brain.

Our experimental design did not employ neuronavigation or a control stimulation site; instead, our participants completed 4 experimental sessions across 4 different days to maximize the variability in the pre-stimulation period to identify stimulation effects robust to state differences (boredom, fatigue, mindwandering, etc). However, an interesting avenue for future work would be to examine network changes that may be more closely tied to functionally-localized regions, where neuronavigation would serve a critical role in equating more precise stimulation locations between individuals. Similarly, our analysis did not require a control site since our investigation examined changes between baseline activity and activity following stimulation. The debate on how to “control” for neurostimulation techniques has recently received increased attention. Research using simultaneous TMS-EEG studies often have non-stimulation related evoked activity (i.e., the auditory “click”) (Conde et al., 2019); however, there is also a debate to how the researchers implemented their controls (Belardinelli et al., 2019). This debate is essential for studies that directly examine the neural activity following stimulation; however, we overcame the challenge of non-stimulation related evoked activity by comparing conditions where these nuisance signals are nearly equated. Thus, our investigation examined changes between baseline activity and activity following stimulation and presents an alternative to this debate within the context of two stimulation sites. The analytic logic we employed in our analysis follows the classic comparison logic between conditions in traditional neuroimaging analysis: conditions are designed to only manipulate the factor of interest, so looking at their difference eliminates all of the concomitant neural processing that occurs but is tangential to condition comparison of interest. Here, the analysis examined differences between occipital and parietal stimulation, so each served as the other’s control for nuisance signals that are concurrent but tangential to the stimulation effects. A similar logic applies to the experimental design decision to not include an explicit control for the auditory click sound from the TMS pulse. Although participants wore ear plugs to mitigate the sensory response in auditory cortex, the focus on relative differences between stimulation sites should help eliminate the effect of the auditory response on the results since it is expected that the sensory effect is equivalent across the conditions. However, future research could examine any of these assumptions in greater detail. In particular, investigations may use more regions and intensities to augment our understanding about differences in ongoing activity with more functionally-determined stimulation protocols.

### 5.5 Conclusion

Using a recently developed network-based methodology applied to EEG, we have investigated the reconfigurations of naturally resonating communities of brain regions. While the alpha network reveals the dynamic interplay of local and global activity, communities within the beta band revealed a remarkable specificity, displaying more local connectivity changes. Particularly important next steps include linking these observations with emerging theories of the impact of stimulation on distributed networks in the form of network control theory (Gu et al., 2015), which has begun to offer insights into the brain’s preference for certain low energy states (Betzel et al., 2017), the role that brain topology plays on the energy requires for brain state transitions (Kim et al., in press; Tang and Bassett, in press), and the predicted impact of stimulation on distal areas (Gu et al., in press; Muldoon et al., 2016). Efforts to ground TMS studies like the one we report here in a mechanistic theory could have lasting consequences in studies linking behavioral changes to neural oscillations or neurostimulation (Medaglia et al., in press), but may also impact future protocols for clinical purposes, providing a means to reconfigure resonating clusters in the brain.

## 6 Materials and Methods

### 6.1 Participants

Ten individuals (7 men, 3 women, aged 20-33, M = 23.8, STD = 4.8) participated in the TMS-EEG experiment. All gave informed, written consent as approved by the University of California, Irvine Institutional Review Board.

### 6.2 TMS/EEG Data Collection

Data collection occurred in the TMS/EEG Laboratory in the Human Neuroscience Lab at the University of California, Irvine. The subjects were seated in a comfortable chair approximately 60 cm from the monitor, equipped with earplugs to attenuate the sound of the coil discharge, and their heads were fixed in a chinrest to minimize movement while they continuously fixated on the center of the monitor screen. No overt motor responses were required during the 30 min experimental session.

Stimulation was applied with a MagStim Model 200 Monophasic Stimulator P/N 3010-00 equipped with a figure-of-eight coil at 55% stimulator intensity (E-field 297 V/m). We established this intensity in a previous study which had a similar protocol (Garcia et al., 2011) by systematically modulating the intensity threshold since motor thresholds are inappropriate for occipital stimulation (Stewart et al., 2001). We found that only 37% of participants saw a phosphene at approximatey 70%. Thus, we set the stimulation intensity for this study to be at 55% of stimulation (20% less than the phosphene induction threshold). We targeted four regions that included symmetric areas in occipital and parietal cortices (Figure 6), and the location for these regions were estimated by electrodes O1/O2 and P1/P2 of the 10-20 international scheme for EEG. This method reliably targets a similar area across participants within 2cm of sulcal/gyral landmarks (Herwig et al., 2003), the resolution of high-density EEG used in this study. For occipital stimulation, the coil was oriented parallel to the coronal view (i.e., paddle pointed up), and for parietal stimulation, the coil was oriented at approximately tangential to the curvature of the scalp at the electrode target locations (i.e., paddle pointed back and down). For each region, participants completed a block of approximately 100 single pulses of stimulation that were no closer than 4 sec apart (jittered up to 6 seconds apart), following standard safety procedures (Rossi et al., 2009). Within a session, the 10 blocks were semi-randomly selected among the four stimulation sites (O1, O2, P1, P2), ensuring that multiple blocks of each stimulation type occurred in each session. Participants completed 4 sessions on separate days, providing an aggregate of 4000 total stimulation trials for each subject.

**Figure 6:**
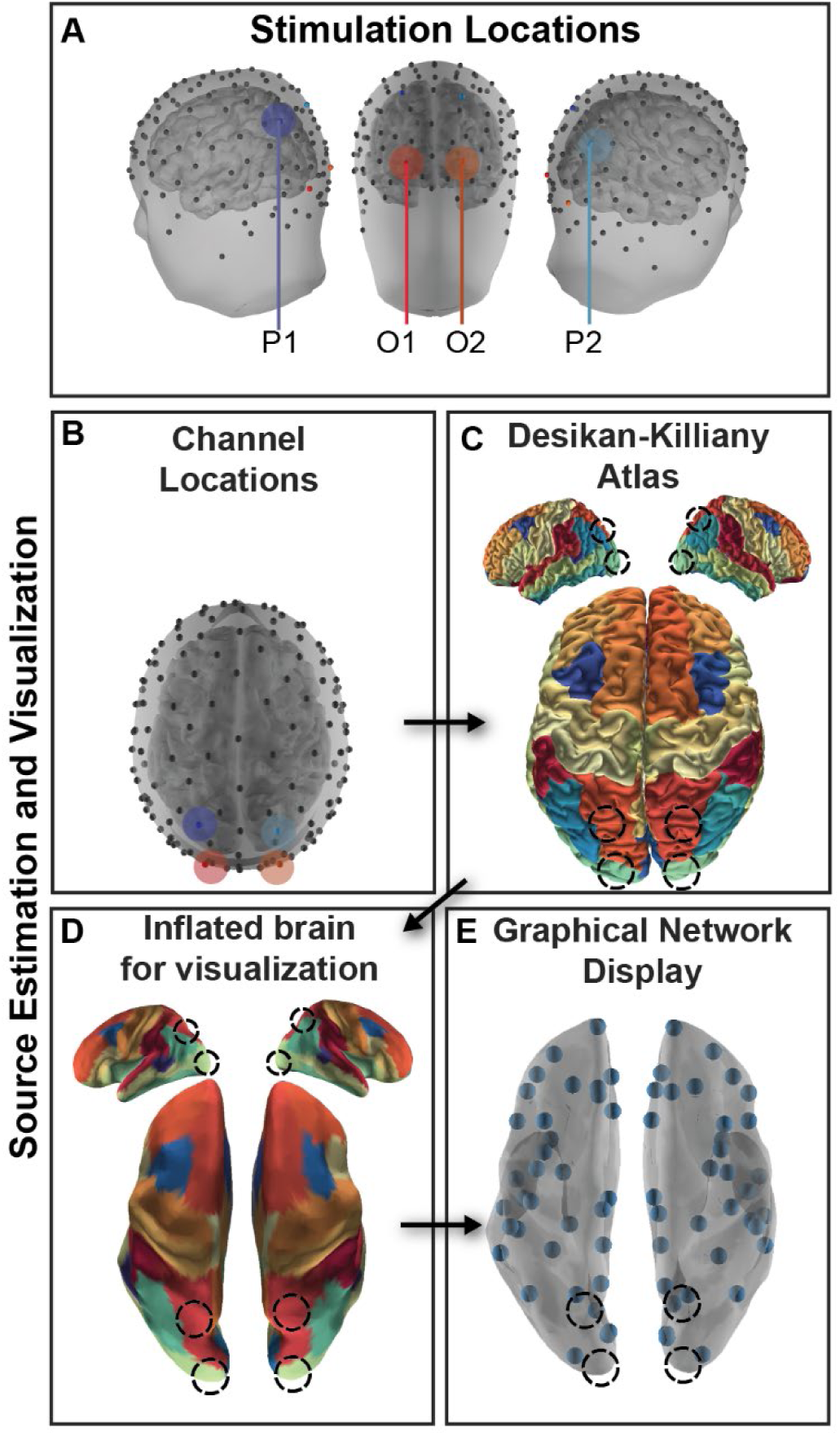
Experimental Design and Analysis. A. Participants received stimulation in symmetric regions in occipital (O1, O2) and parietal (P1, P2) cortex. B. High density EEG recorded from 128 channels was submitted to a cLORETA source analysis, and current source density (CSD) was estimated for each vertex of a high resolution mesh. C. CSD was then averaged within a parcellation of cortex following the Desikan-Killiany atlas parcellation to estimate regional brain activity. D. The cortex was inflated for visualization. E. Each centroid of the region is plotted as a small orb. Stimulation locations marked in each visualization with a shaded region or a dotted line.

**Figure 7:**
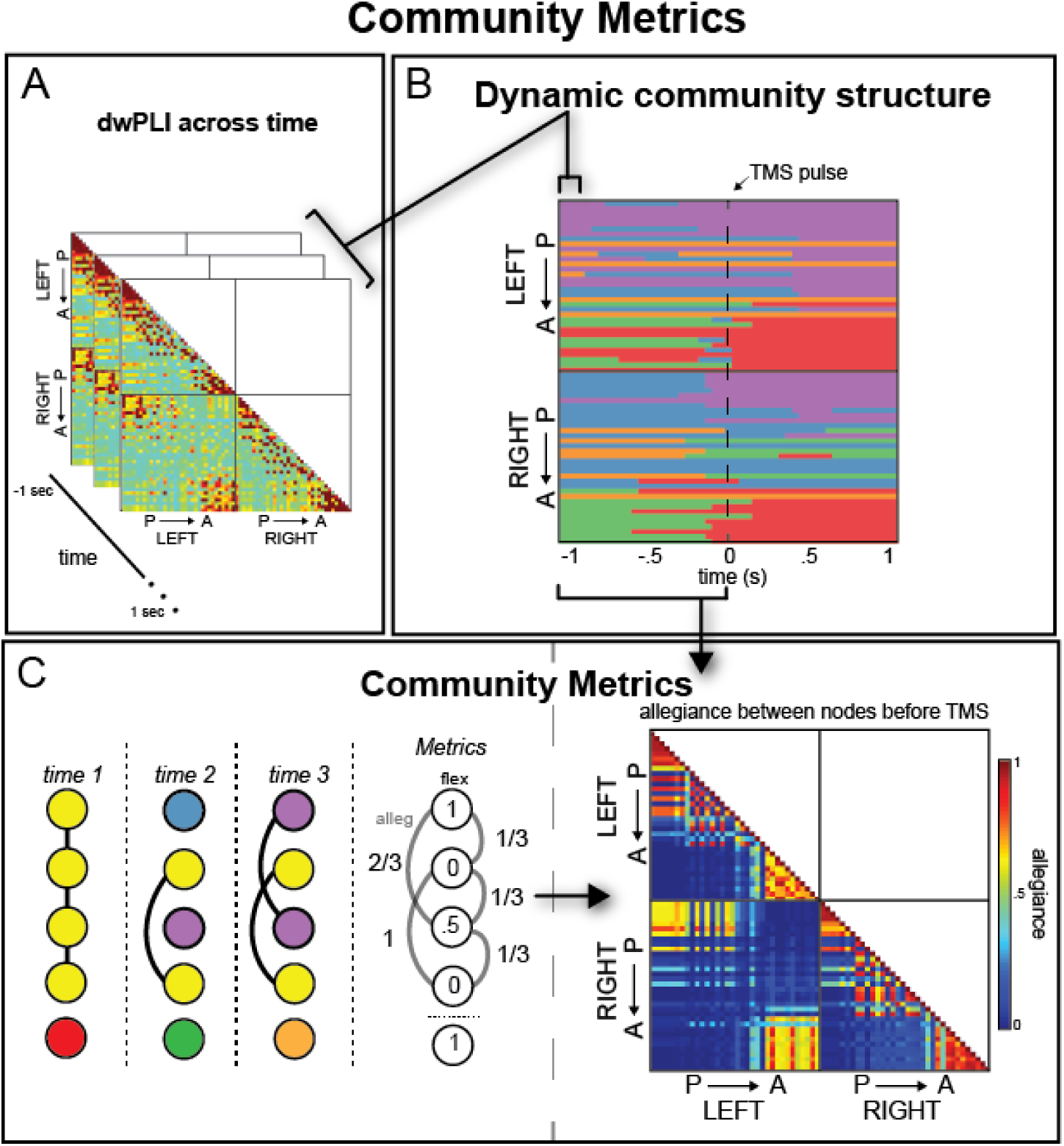
Overview of Analysis method and natural brain architecture. (A) DwPLI was estimated across trials in 40ms windows across the 2sec epoch, including one second before and one second after the TMS onset (P: posterior, A: anterior nodes). (B) Dynamic network communities from a sample subject derived from the dwPLI estimate for each window. Each color marks a different community label. (C, left) From B, community metrics were calculated that represent how often a node changes over time (flexibility) or how often each node-pair is in the same community over time (allegiance). The cartoon networks show 5 hypothetical nodes that change communities over time (3 time windows shown). Connections are marked as black lines and metrics are given in the final column, showing a range of flexibility and allegiance values. These metrics are used in subsequent analyses. (C, right) Summary allegiance matrix for a sample subject for the period before TMS onset, indicating the natural architecture of connectivity within the alpha band.

Throughout the session, 128 channels of EEG were recorded using a TMS-compatible EEG system from Advanced Neuro Technology (ANT, The Netherlands). The Waveguard cap system consisted of small Ag/AgCl electrode elements, specially designed to yield high-quality and stable recordings with simultaneous TMS. EEG data were sampled at 1024 Hz for this experiment and impedances were targeted to be kept below 10 kΩ.

### 6.3 EEG Analysis

#### 6.3.1 Segmentation of epochs

The EEG data were preprocessed following an established pipeline (Garcia et al., 2011) for each of the four sessions for each of the ten participants (40 total datasets). First, raw EEG were submitted to a principal component analysis (PCA) to identify the peak amplitude events that correspond to the stimulation event and to eliminate any timing discrepancy between the intended and actual timing of the stimulation pulse in the EEG data. After the timing of each pulse was recovered, the decomposition was discarded and was not used in any subsequent analysis (i.e., no components were removed). Using the maxima from the largest component of each dataset as the onset of stimulation, the 40 datasets were segmented into 40,000 epochs that were two seconds in duration, including one second (1024 samples) before and after the TMS pulse. After this segmentation procedure, the 4 samples before the pulse and 16 (15.6 ms) after the pulse were removed from the epochs to remove artifact from the amplifier. These samples were later replaced with a forward autoregressive moving average prediction of the contaminated data from the intervals directly preceding the TMS pulse and a mild Savitzky-Golay smoothing filter over the interval surrounding the pulse to remove any quick shifts in amplitude due to the artifact editing procedure. Finally, each trial was normalized by the standard deviation of the 512 samples prior to the stimulation. Due to the known multiple sources of artifact in work using simultaneous TMS-EEG (Rogasch and Fitzgerald, 2013) beyond the amplifier artifact within the 20 samples (19.5ms), aggressive means were used to ensure artifact was eliminated from our estimates. Automated artifact editing based on amplitude thresholds was used to eliminate approximately 30% of trials likely contaminated with artifacts from the TMS pulse or blink artifacts, leaving the following trial total for analysis in the participants: 3,129, 1,768, 3,535, 4,021, 2,269, 2,308, 2,931, 2,032, 2,840, and 3,088. After this preliminary pre-processing, we largely followed the PREP approach, an artifact removal procedure that has been shown to be robust to artifacts within environments high artifact environments (Bigdely-Shamlo et al., 2015). The following steps were completed (1) line noise removal via a frequency-domain (multi-taper) regression technique to remove 60Hz and harmonics present in the signal, (2) a robust average reference with a Huber mean, (3) artifact subspace reconstruction to remove residual artifact with the standard deviation cutoff parameter was set to 15, and (4) band-pass filtering using a Butterworth filter with 2-dB attenuation at 2 and 50 Hz. TMS-evoked potentials (TEPs) may be seen in Supplemental Figures 2, 4, and 5, displaying how the preprocessing pipeline has successfully removed this data.

#### 6.3.2 Distributed Source Reconstruction

From the pre-processed EEG data, we estimated current source density (CSD) over a 5003-vertex cortical mesh. A boundary element method (BEM) forward model was derived from the ‘Colin 27’ anatomy (Holmes et al., 1998) and transformed into MNI305 space (Evans et al., 1993) using standard electrode positions fit to the Colin 27 head surface in BrainStorm (Tadel et al., 2011). The BEM solution was computed using OpenMEEG (Kybic et al., 2005; Gramfort et al., 2010), and the cLORETA approach was used for inverse modeling as described in detail in (Mullen et al., 2015) and implemented in the BCILAB (Kothe and Makeig, 2013) and Source Information Flow (Mullen, 2014) toolboxes. Then CSD was averaged into one time course from each of the 68 regions of the Desikan-Killiany atlas (Desikan et al., 2006) and down sampled to 128Hz for use in the connectivity analysis. As a final step after parcellation, the surface mesh of the Colin 27 brain (with DK atlas parcellated regions) was imported into Matlab and distance was estimated between regions closest to the stimulation site (bilateral lateral occipital and superior parietal) and every other region in the DK atlas using pdist.m in Matlab. For community ‘distance’, the average distance from the stimulation site was estimated, corresponding to centroid distance of the community.

### 6.4 Functional Connectivity Analysis

To estimate functional connectivity of brain networks before and after stimulation, an undirected measure, known as the debiased weighted phase lag index (dwPLI), was computed between each pair of the 68 estimated brain regions from the atlas. DwPLI is robust against the influence of volume conduction, uncorrelated noise, and intersubject variations in sample size (Vinck et al., 2011; Vindiola et al., 2014), and it has previously been proposed to be an appropriate pairing with a source localization analysis to minimize the influence of these nuisance variables (Hillebrand et al., 2012; van Diessen et al., 2015).

The connectivity estimates were calculated using Matlab and FieldTrip (Oostenveld et al., 2010). First, a multitaper spectral estimation was applied to the CSD measurements, and then dwPLI was computed between all source pairs to estimate the functional connectivity pattern of signals in the frequency range between 2Hz and 25Hz (step = 0.5 Hz) in a 5 sec window centered on the stimulation pulse (−2.5sec to 2.5sec). The dwPLI connectivity estimates were calculated across trials, representing the trial-by-trial consistency between regional CSD. Next the matrix of dwPLI estimates was reduced to the 51 time windows corresponding to windows centered 38ms apart, from 1 sec before the TMS pulse to 1 sec after the TMS pulse. Since this estimate was done using a hanning windowing method, the windows are not independent and represent some smearing in time.

### 6.5 Dynamic Community Detection

In addition to looking at whole-brain connectivity patterns, we employed a community detection algorithm (Bassett and Bullmore, 2006; Bullmore and Sporns, 2012; Ercsey-Ravasz et al., 2013) to examine whether regions formed modular networks, and if the regional composition of these networks changed before and after stimulation across the 51 time windows in our 2 second epoch. The algorithm optimizes a multilayer modularity quality function, Q, using a Louvain-like greedy algorithm (Blondel et al., 2008; Mucha et al., 2010) to assign brain regions to communities. The community assignments are dependent on two parameters: (1) a structural resolution γ parameter and (2) a temporal resolution ω parameter. These two parameters determine the scale of the resulting graph, both structurally and temporally, and here, we sweep this parameter space to find the scale of the data that is most unlike that expected in an appropriate random network null model. As described in Garcia et al. (2018), there are several heuristics we may use to determine the optimal parameter for our dataset. We chose an unbiased “difference” heuristic because of the unique properties of this stimulation dataset, which we explain below.

Following our previous work on fMRI data (Bassett et al., 2013), the values for both parameters were determined by comparing the mean value of Q in the experimental data to the mean value of Q in a shuffled null model of the data; we tested a very wide range of values for each parameter since this algorithm has not yet been applied to EEG data, which has inherently different temporal and spatial scales of functional connectivity (Nunez and Srinivasan, 2006). Our analysis examined parameters for γ = 0.8 to γ = 1.6 and ω = 0.5 to ω = 35. The null model of the data was created by randomly shuffling the pair-wise dwPLI values, destroying the correlational structure observed in EEG data for each subject and parameter pairing. Each Q was then subtracted for each parameter pairing, comparing the observed model’s Q (from the unperturbed EEG connectivity patterns) and the null model’s Qnull (shuffled connectivity patterns) for each subject, and our analysis found a clear peak in the resulting Q matrix, suggesting that the range used was appropriate for this dataset. In fact, the largest difference was found for γ = 1.025 and ω = 9, and these parameters were used in the reported analyses, which suggests that the temporal parameter (ω) is the parameter that captures the unique properties of the EEG signal. Since the community detection algorithm is non-deterministic (Good et al., 2010), 100 iterations of the hard partitions were estimated with modularity maximization for each subject and stimulation condition (O1, O2, P1, P2), yielding 100 sets of community labels for the 68 nodes for each of the 4 stimulation conditions for each of the 10 subjects.

#### 6.5.1 Community Metrics

Within each of the 51 time windows of our 2 second stimulation epoch, we examined the relationship among the brain regions within a community to characterize the dynamic reconfiguration of spatially distributed neural sources before and after stimulation. Our analysis investigated two community metrics, flexibility (Bassett et al., 2011) and allegiance (Bassett et al., 2015).

The *flexibility* of each node corresponds to the number of instances in which a node changes community affiliation, g, normalized by the total possible number of changes that could occur across the layers L (Bassett et al., 2011), which represents each time slice within this dynamic community detection algorithm. In other words, the flexibility of a single node *i*, *ξ_i_*, may be estimated with:

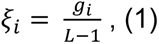

where L is the total number of temporal windows.

*Allegiance* estimates how much regions communicate with subnetworks in the community structure and demonstrate the same pattern of connectivity across time points. We define allegiance matrix P, where edge weight Pij denotes the number of times a pair of nodes moves to the same community together divided by L−1 possible changes.

Thus, allegiance increases the resolution of community and captures coordinated activity of each node with every other node in the brain whereas flexibility examines *whether* a brain region changes affiliations overall.

Each of these measures was calculated twice, once for the 25 windows of partitions before TMS (preTMS) and once for the 25 windows following TMS (postTMS), ignoring the center window where stimulation occurred. Our analysis focused on the absolute difference in allegiance and flexibility between preTMS and postTMS communities, emphasizing whether stimulation influences local or global brain dynamics more strongly.

Finally, results were averaged across left and right stimulation sites since we were interested in the general magnitude of changes from functionally similar regions. In support of this data reduction primarily driven by our broad interest in coarse parietal/occipital stimulation differences, follow-on analyses did not show any differences in consistency of network changes between left and right stimulation, using a temporal consensus method inspired by Doron and colleagues in Supplemental Figure 8 (Doron et al., 2012).

### 6.6 Statistical Analyses

To find the substantial changes in metrics across the pre-TMS and post-TMS intervals, traditional linear statistics were used, where the pre-TMS and post-TMS intervals were treated as *conditions* and paired-sample t-tests were applied to node or communities as indicated in the text. In cases where multiple comparisons were carried out (e.g., Figure 3), a Bonferroni correction to the alpha value was used to determine significance. For the correction, each band-specific metric was treated as a separate set of tests (10 comparisons within a set, Community 1 vs Community 2, Community 1 vs Community 3, etc), so the corrected alpha value was set to 0.005. Where appropriate, both the corrected and uncorrected significant comparisons are shown (see Figures 3 and 4).

## 8 Acknowledgments

The authors acknowledge thoughtful discussions with our colleagues at University of California, Irvine for study coordination and subject testing. This research was supported by mission funding to the Army Research Laboratory as well as sponsored by the Army Research Laboratory and accomplished under Cooperative Agreement Number W911NF-10-2-0022. We would also like to acknowledge that the work was partially collected while JOG was funded by a National Research Service Award (F31-EY-019241) awarded by NIH. The views and conclusions contained in this document are those of the authors and should not be interpreted as representing the official policies, either expressed or implied, of the Army Research Laboratory or the U.S. Government.

## 9 Supplementary Material

**Supplemental Figure 1:**
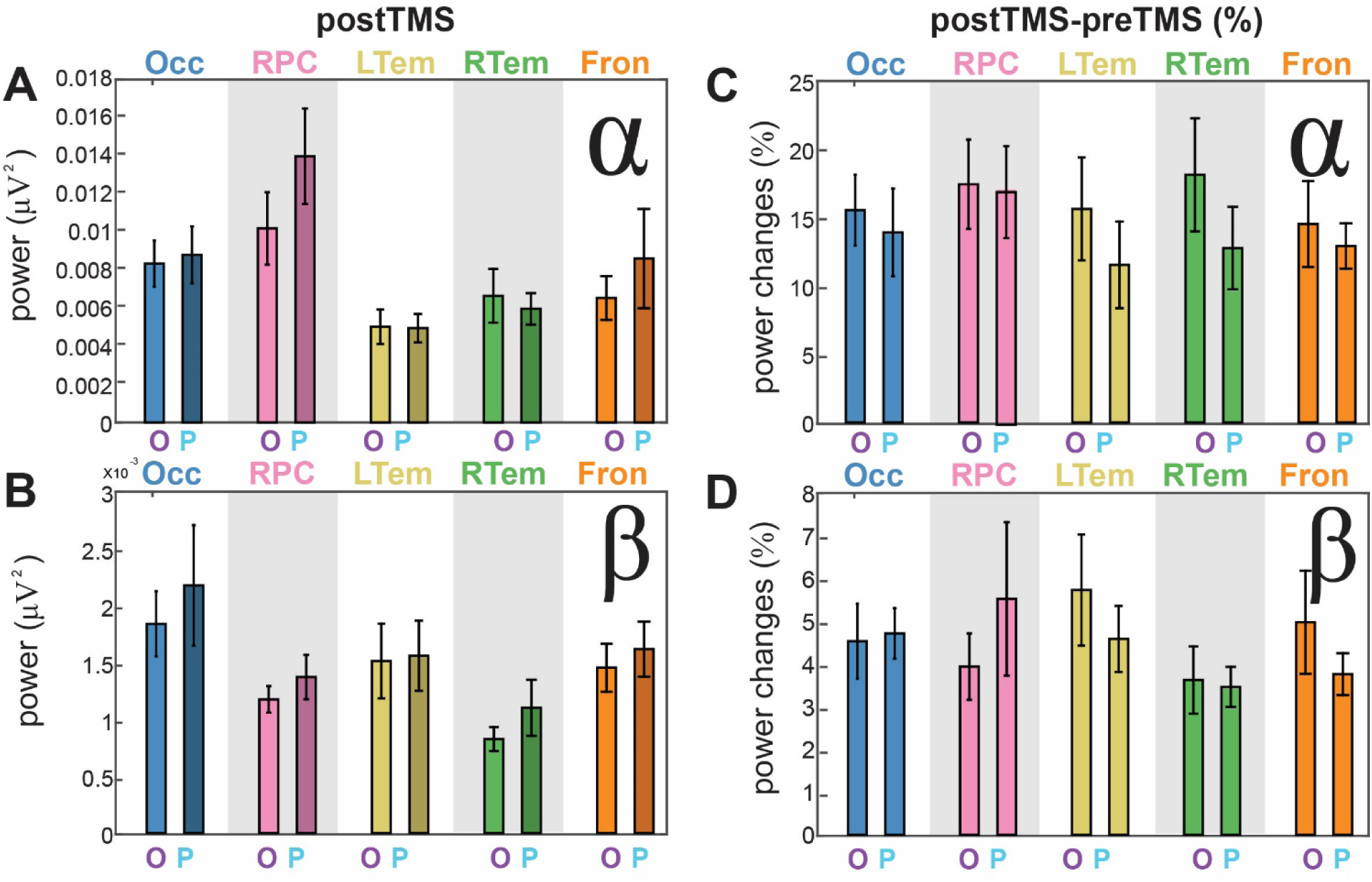
Average power and power changes in communities for occipital (O) and parietal (P) stimulation for the alpha band (A,C) and beta band (B,D). (A,B) Power estimated from the second following stimulation and averaged across nodes within a community and then across participants Error bars indicate the SEM across participants. (C,D) Power changes plotted as percent change from the preTMS level of power. Error bars indicate the SEM across participants

**Supplemental Figure 2:**
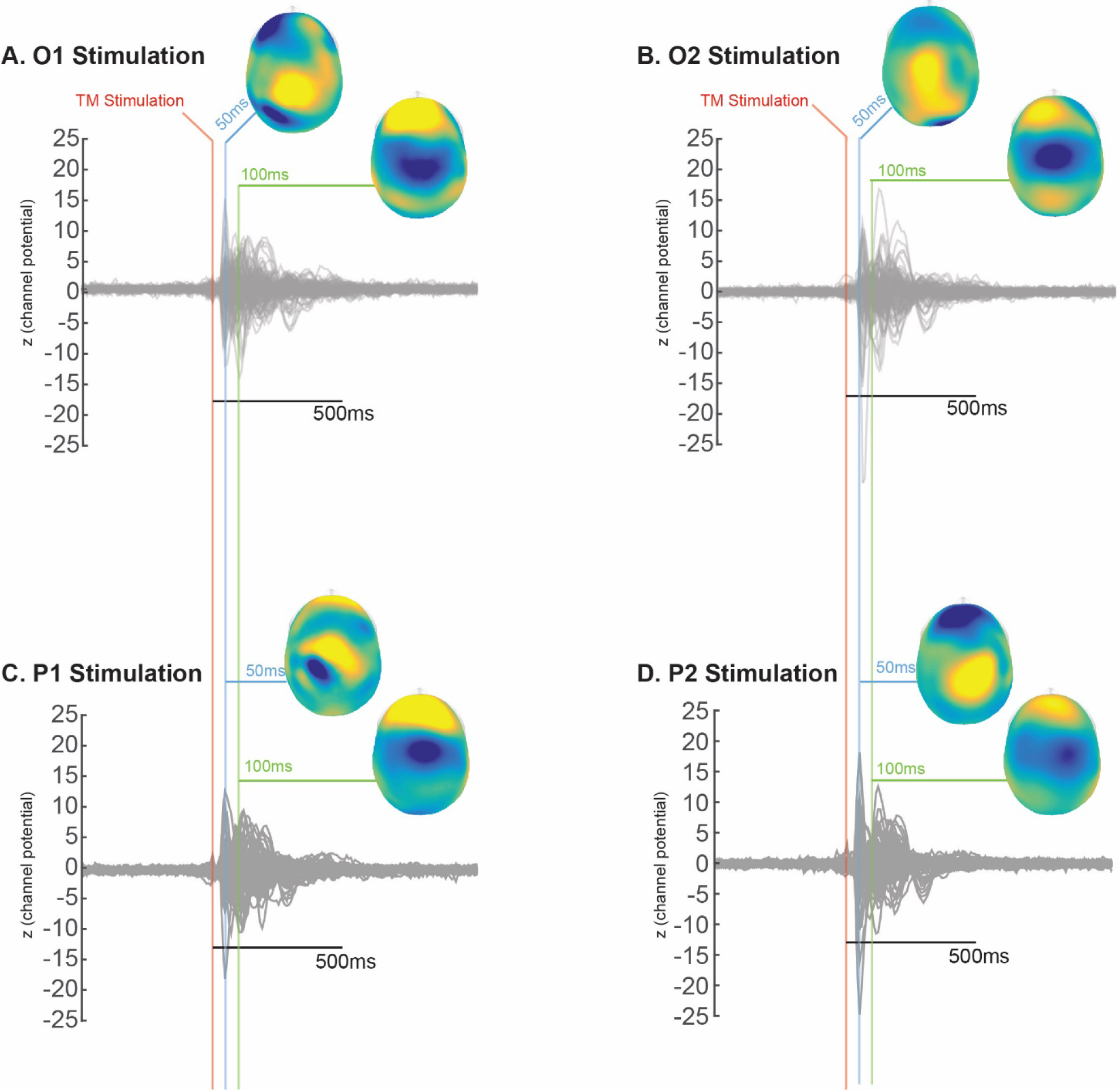
Mean channel traces (normalized by the mean and standard deviation of 500ms of samples before TM stimulation) across participants with topographic plot insets visualizing the amplitude across all 128 channels showing increases (yellow) and decreases (blue) across the scalp. (A,B) Mean evoked response to left (A) and right (B) occipital stimulation show clear and obvious changes near the stimulation site 50ms after stimulation (blue line) and a subsequent general negative deflection 100 ms after stimulation, typical of single pulse TEPs. (C,D) Mean evoked response to left (C) and right (D) parietal stimulation display a similar pattern to occipital stimulation, with a spatial shift anterior to the peaks shown with occipital stimulation.

**Supplemental Figure 3:**
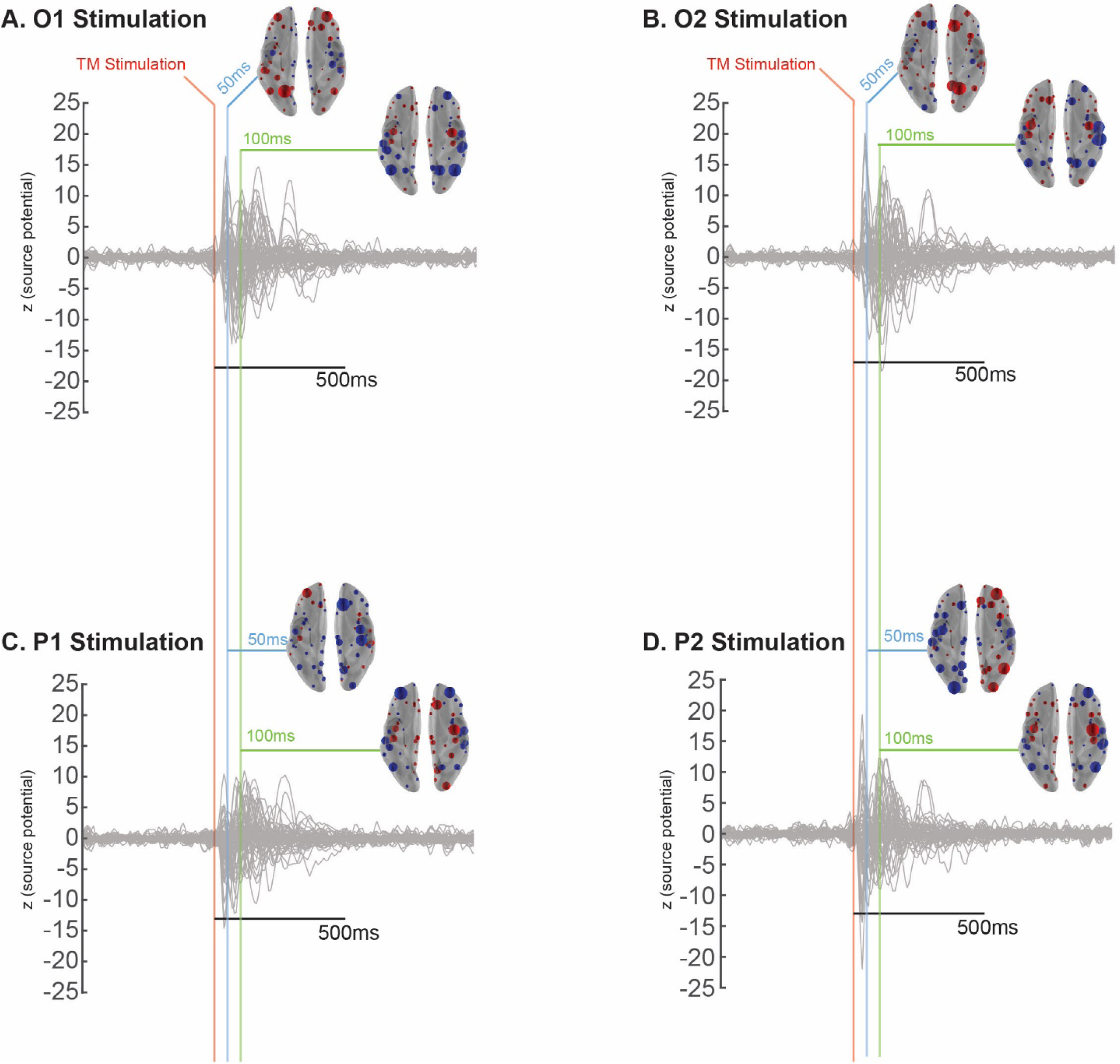
Mean source potentials (normalized by the mean and standard deviation of 500ms of samples before TM stimulation across participants with brain insets visualizing the source potential across all 68 regions of the brain (scaled within each time slice). (A,B) Mean evoked source potential to left (A) and right (B) occipital stimulation show a clear enhancement near the stimulation site 50ms after stimulation (blue line) and a subsequent reversal 100 ms after stimulation, typical of single pulse TEPs. (C,D) Mean evoked source potential to left (C) and right (D) parietal stimulation display slightly more variability for left stimulation, but the standard enhancement followed by reversal for right stimulation.

**Supplemental Figure 4:**
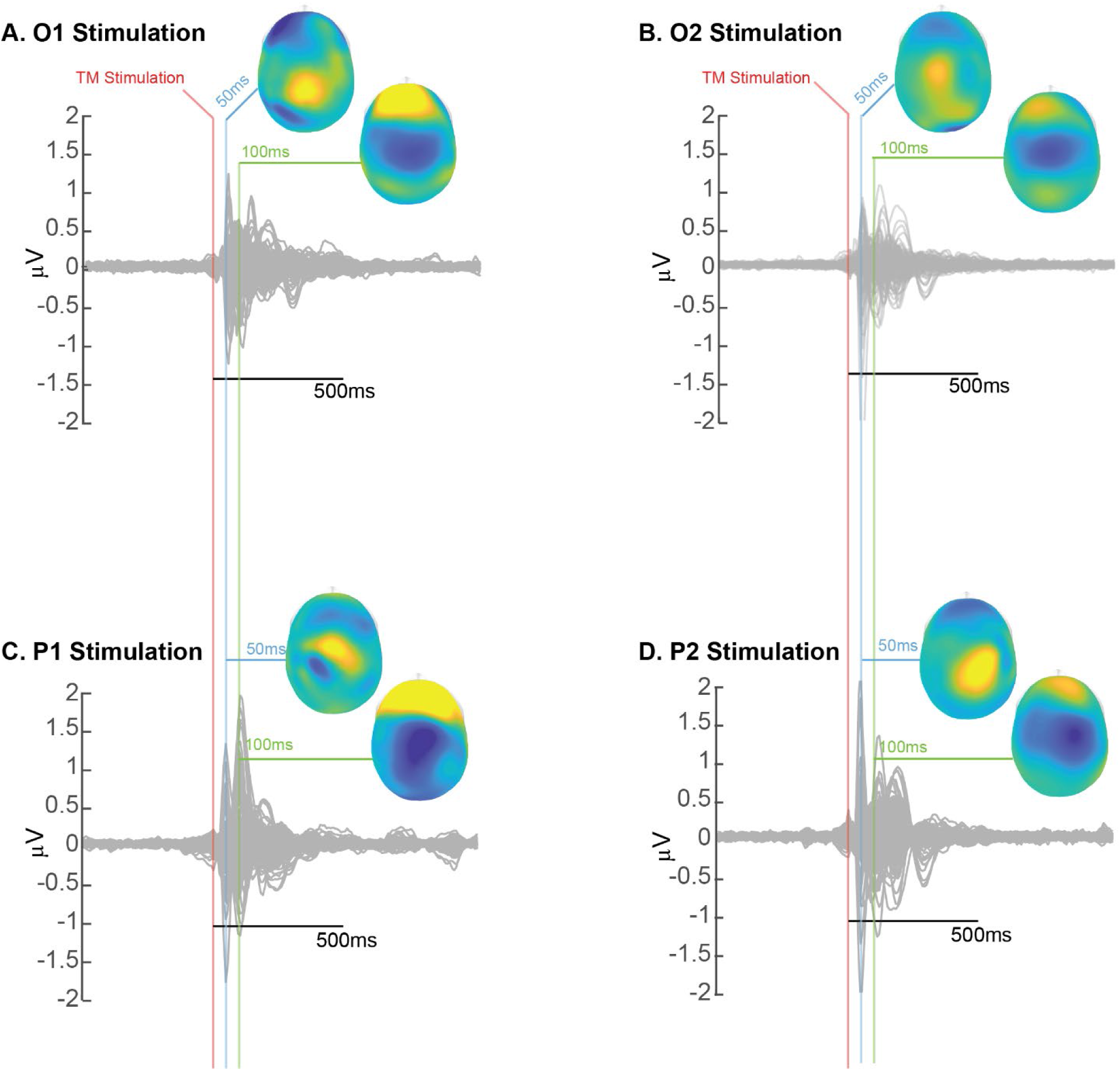
Mean channel traces (with baseline subtraction 500ms of samples before TM stimulation) across participants with topographic plot insets visualizing the amplitude across all 128 channels showing increases (yellow) and decreases (blue) across the scalp. (A,B) Mean evoked response to left (A) and right (B) occipital stimulation show clear and obvious changes near the stimulation site 50ms after stimulation (blue line) and a subsequent general negative deflection 100 ms after stimulation, typical of single pulse TEPs. (C,D) Mean evoked response to left (C) and right (D) parietal stimulation display a similar pattern to occipital stimulation, with a spatial shift anterior to the peaks shown with occipital stimulation.

**Supplemental Figure 5:**
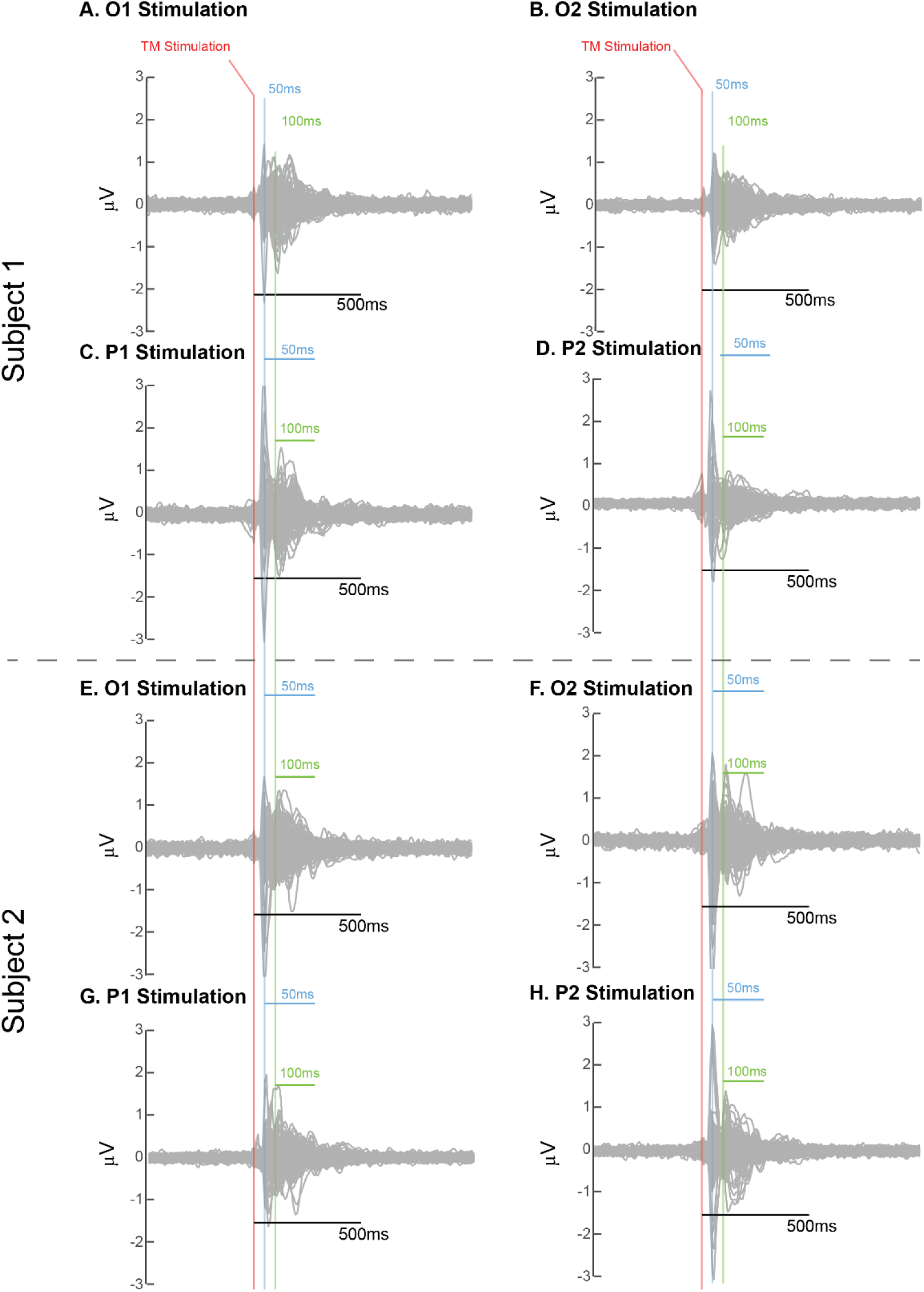
Mean channel traces (with baseline subtraction 500ms of samples before TM stimulation) within two sample subjects, highlighting the consistency in timing of the evoked responses, but also the small changes in amplitude across participants. (A,B,E,F) Mean evoked response to left (A) and right (B) occipital stimulation show clear and obvious changes near the stimulation site 50ms after stimulation (blue line) and a subsequent general negative deflection 100 ms after stimulation, typical of single pulse TEPs. (C,D,G,H) Mean evoked response to left (C) and right (D) parietal stimulation display a similar pattern to occipital stimulation.

**Supplemental Figure 6:**
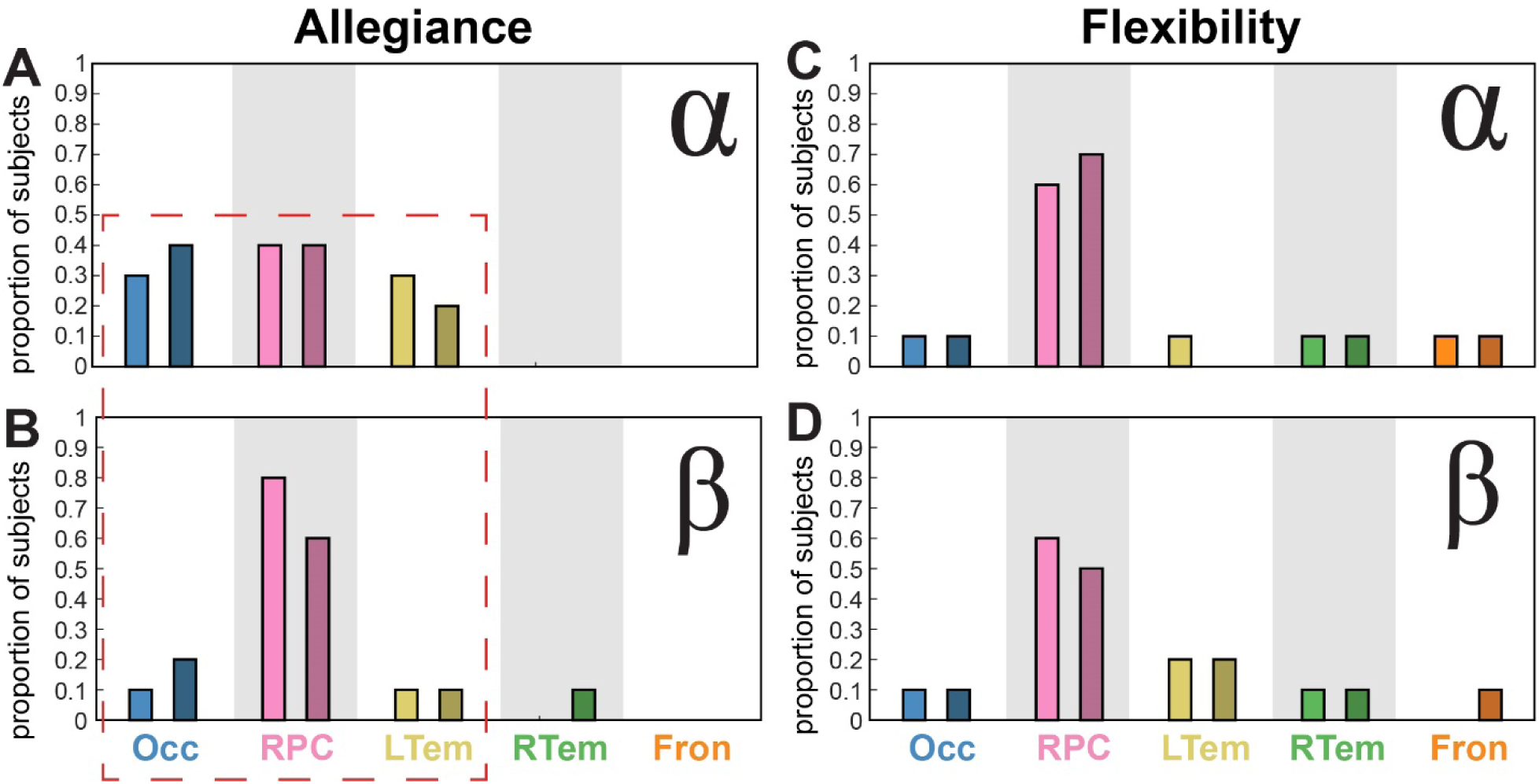
Bar plots show the proportion of subjects who have the highest allegiance (A,B) and flexibility (C,D) in each frequency band, providing some evidence of robustness of the results. Highlighted in red, estimated allegiance in 6 and 8 subjects in the RPC community is the highest in the beta band for occipital and parietal stimulation, respectively. In contrast, estimated allegiance in the 3 communities closest to the stimulation site is similarly robust across these communities.

**Supplemental Figure 7:**
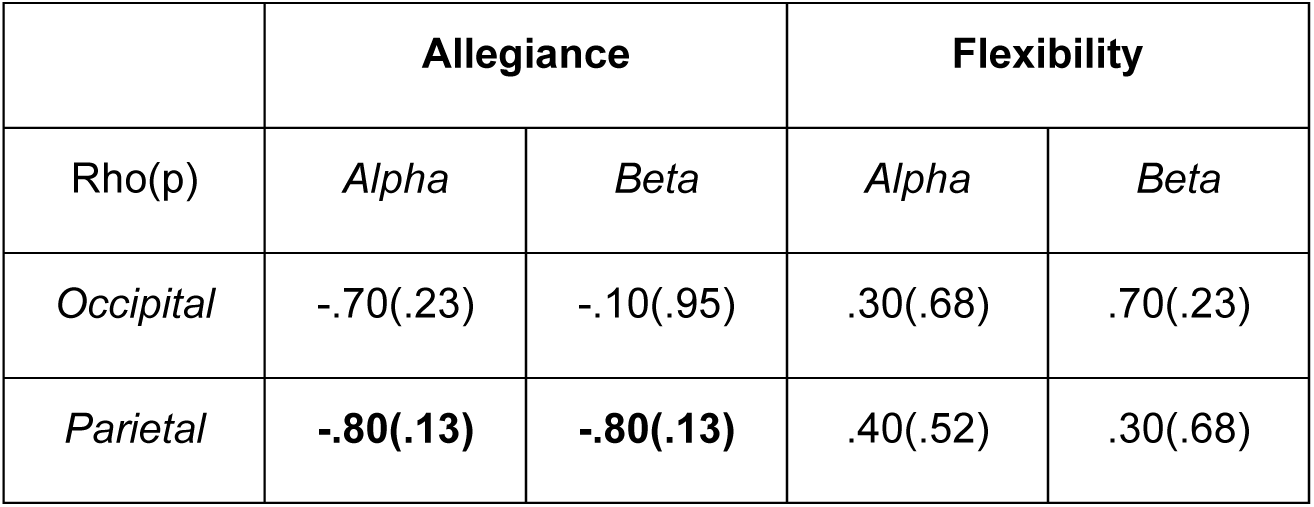
Spearman’s Rho non-parametric correlation analysis correlating Allegiance and Flexibility with distance from the stimulation site. The analysis reveals opposing trends (negative *Rho* for allegiance and positive *Rho* for flexibility), but none of the results were significant (all p > .05). Parietal stimulation showed the strongest relationship, as indicated by the largest magnitude in Rho and the lowest p-value (p = 0.13). This analysis confirms much of what we see in the data; however, the non-significant effects do not change our interpretation.

**Supplemental Figure 8:**
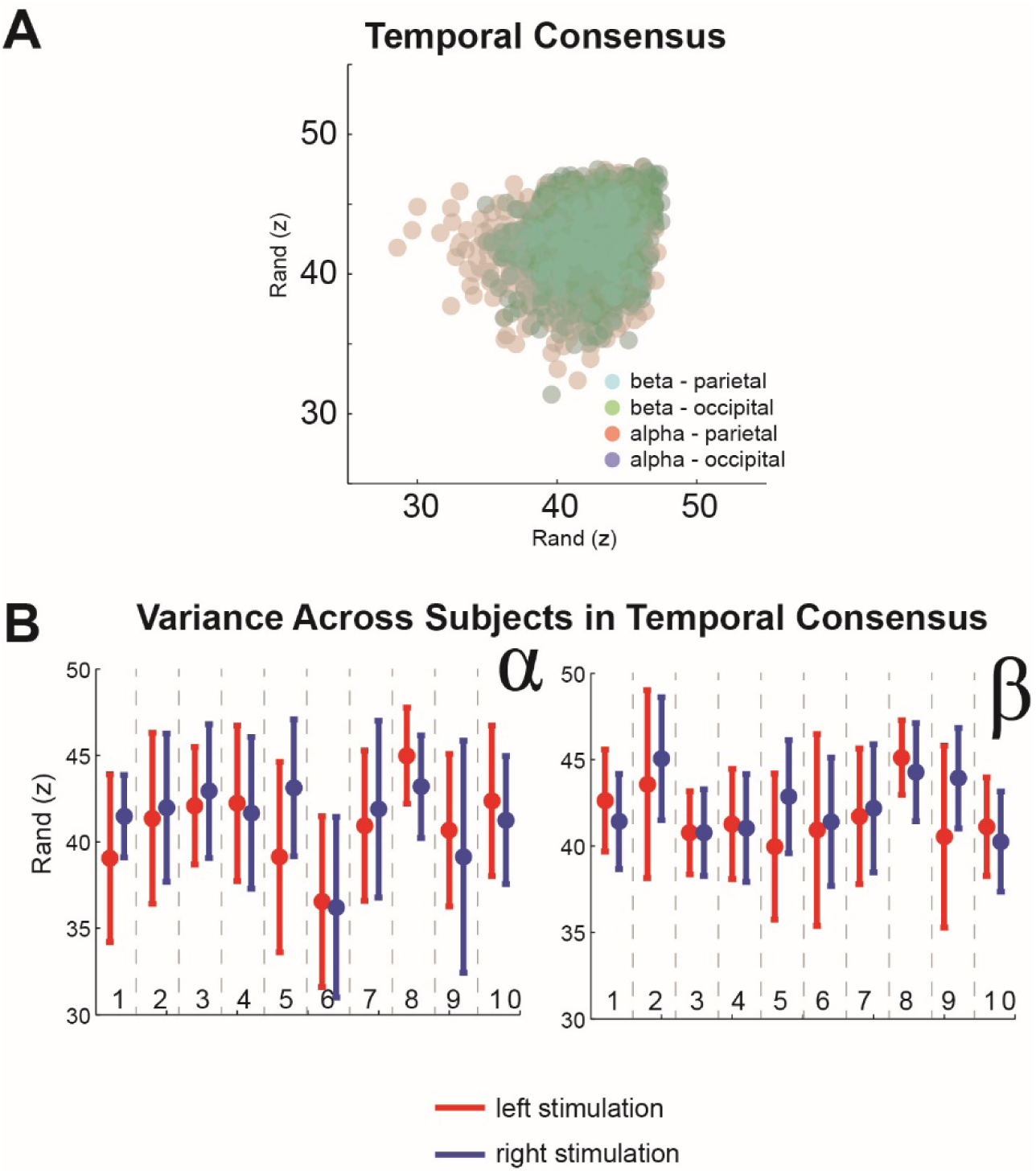
Left and right stimulation similarity found with the Rand index, a metric of similarity between clusters (A) We plot the Rand z-score for each application of the community detection algorithm (100 iterations for each of 10 subjects), where each semi-transparent dot is colored to indicate stimulation condition (parietal or occipital) and frequency band (alpha or beta). Across all participants, only a very narrow range of Rand z-scores is observed, with high variability between all conditions. Due to this difference across conditions and iterations, we examined the variance within each subject across the 100 iterations within each condition. (B) Rand z-scores for alpha are in the left plot and beta in the right plot with scores for stimulation to right hemisphere in blue and left hemisphere in red. The error bars indicate the 95% confidence of the mean estimate. As illustrated in the plot, each subject’s distributions demonstrate narrow overlapping ranges, where between-subject variability is much more substantial than the left-right stimulation difference.

## Notes

#### Summary of Updates

Substantial text revisions and supplemental material additions.

